# Morphological integration and evolutionary potential of the primate shoulder: Variation among taxa and implications for genetic covariances with the basicranium, pelvis, and arm

**DOI:** 10.1101/2022.06.09.495317

**Authors:** Elizabeth R. Agosto, Benjamin M. Auerbach

## Abstract

Within the primate order, the morphology of the shoulder girdle is immensely variable and has been shown to reflect the functional demands of the upper limb. The observed morphological variation among extant primate taxa consequently has been hypothesized to be driven by selection for different functional demands. Evolutionary analyses of the shoulder girdle often assess this anatomical region, and its traits, individually, therefore implicitly assuming independent evolution of the shoulder girdle. However, the primate shoulder girdle has developmental and functional covariances with the basicranium and pelvic girdle that have been shown to potentially influence its evolution. It is unknown whether these relationships are similar or even present across primate taxa, and how they may affect morphological variation among primates. This study evaluates the strength of covariance and evolutionary potential across four anatomical regions: shoulder girdle, basicranium, pelvis, and distal humerus. Measures of morphological integration and evolutionary potential (conditioned covariance, and evolutionary flexibility) are assessed across eight anthropoid primate taxa. Results demonstrate a consistent pattern of morphological constraint within paired anatomical regions across primates. Differences in evolutionary flexibility are observed among primate genera, with humans having the highest evolutionary potential overall. This pattern does not follow functional differences, but rather a separation between monkeys and apes. Therefore, evolutionary hypotheses of primate shoulder girdle morphological variation that evaluate functional demands alone may not account for the effect of these relationships. Collectively, our findings suggest differences in genetic covariance among anatomical regions may have contributed to the observable morphological variation among taxa.

## 1. Introduction

The primate shoulder girdle exhibits the greatest interspecific morphological variation of all placental mammal orders (Oxnard, 1968; Preuschoft et al., 2010). The observed morphological variation among primates matches the diversity of upper limb functional demands, and corresponds to the size, shape, and orientation of the muscles attaching to the bony elements of the shoulder girdle, especially the scapula (Ashton and Oxnard, 1964; Oxnard, 1967, 1968; Badoux, 1974; Roberts, 1974; Ashton et al., 1976; Larson and Stern, 1986, 1992; Larson, 1993, 1995, 2015; Irwin and Larson, 2000; Young, 2008; Preuschoft et al., 2010). Morphological differences that correspond with functional variation in the scapula are present from birth and are maintained through ontogeny (Young, 2006). This indicates that variation in scapular shape is not solely a plastic response to biomechanical demands encountered during the lifetime of individuals. Nevertheless, whether this variation arose in response to direct selection brought about through physical demands on the shoulder related to its function, by correlated responses to directional selection on integrated traits (e.g., Rolian et al., 2010; Rolian, 2014; Grabowski and Roseman, 2015), or through neutral processes remains undetermined.

Studies of morphological variation in the primate shoulder girdle are often focused solely on the scapula (Schultz, 1930,1950; Ashton and Oxnard, 1961, 1964a, b; Oxnard, 1963,1967, 1968, 1969; Ashton et al., 1965; Roberts, 1974; Shea, 1986; Inouye and Shea, 1997; Taylor, 1997; MacLatchy et al., 2000; Taylor and Slice, 2005; Larson et al., 2007; Green, 2013; Green et al., 2106; Feuerriegel et al., 2017; Selby and Lovejoy, 2017), as it serves as a main attachment point for many of the muscles associated with upper limb and shoulder movement. This muscle-bone relationship informs a form-function model where the form of the scapula reflects movements of the shoulder region. Using this model, researchers have drawn evolutionary conclusions in their studies that implicitly make two assumptions: 1) trait differences among taxa result from directional selection acting on those morphologies; and 2) variation within individual traits between primate species reflect adaptations for specific motor functions or environments. While these conclusions may be merited, most studies have not used evolutionary models to quantify how evolution could have occurred, or might occur in the future, to explain interspecific variation (cf. Roseman and Auerbach, 2015). Rather they use a framework based on comparative anatomy. Moreover, the individual assessment of traits implicitly assumes their independent evolution, an assumption that is not supported by the evolutionary biology literature for most traits (Lande, 1979; Lande and Arnold, 1983; Arnold, 1983; Ackermann and Cheverud, 2000; Hansen and Houle, 2008; Walsh and Blows, 2009; Rolian et al., 2010; Roseman and Auerbach, 2015; Katz et al., 2016; Savell et al., 2016). Traits within an organism do not exist in isolation. Rather, they covary with whole-organismal traits (such as body size) and with traits from other anatomical regions that share mechanical and developmental variation.

In a recent study, Agosto and Auerbach (2021) used a quantitative genetics approach to assess the potential evolutionary covariance between the shoulder girdle and two anatomical regions with which it shares mechanical or developmental covariance: the basicranium and pelvic girdle. A novel result of this study indicates that traits of the basicranium and pelvic girdle can exert similar degrees of evolutionary constraint on traits of the shoulder girdle. Furthermore, the results suggest the basicranium imposes evolutionary constraint on both the shoulder and pelvic girdles but is not equally constrained by either the shoulder girdle or pelvic girdle. Additional analyses by Agosto and Auerbach (2021), which include traits of the proximal humerus, also demonstrate greater potential evolutionary independence of these traits from the basicranium and pelvis compared with the scapula. This result aligns with known developmental and functional relationships among these anatomical regions. Overall, Agosto and Auerbach (2021) demonstrated that the evolution of the primate shoulder is more complex than previously perceived, uncovering evolutionary covariance with anatomical regions once assumed to be independent of the shoulder (i.e., the basicranium). These relationships, however, are not evident when evaluating the scapula or its traits alone.

While the study by Agosto and Auerbach (2021) established a model where the evolution of the shoulder girdle is influenced by its underlying covariances with anatomical regions in which it shares functional and developmental relationships, a non-comparative approach was taken by investigating these relationships within a single genus: *Colobus*. How these relationships relate to the larger breadth of primate shoulder girdle morphological variation remains unanswered. Researchers cannot assume that the results indicated for the monkey taxa investigated by Agosto and Auerbach (2021) are representative of the potential evolutionary relationships among the three anatomical regions for apes, let alone all primates. For example, other studies (Young, 2004; Rolian, 2009; Young et al., 2010; Lewton, 2012) have demonstrated that integration, and thus the evolvability, of postcranial traits vary considerably among primates, and that apes generally are less integrated than monkeys. The current study therefore expands upon the findings of Agosto and Auerbach (2021) by comparing the patterns of evolutionary potential among anatomical regions between a broad sample of primate genera.

### 1.1. Study predictions

By investigating morphological integration and estimates of evolvability, we seek to understand how trait covariance that has resulted from developmental and mechanical relationships between the shoulder, basicranium, and pelvis varies among primate genera, and whether these could have influenced the observed diversity in shoulder girdle morphology among primates. We use a model where shared development and function influence evolutionary potential among these anatomical regions. Additionally, we include measures of the distal humerus as a more independent anatomical region under our model, as the distal humerus has no direct muscle attachments or known shared developmental processes or factors with either the basicranium or pelvis. We asked the following two questions:

1. Is there evidence that evolutionary constraint among the basicranium, shoulder girdle, and pelvic girdle is present amongst a range of primate taxa?
2. Do patterns of evolutionary constraint among primates correspond to functional variation of the upper limb as broadly reflected by locomotion (e.g., Ashton et al., 1976; Roberts, 1974; Larson, 1993, 1995; Irwin and Larson 2000)?

To address these questions, the magnitude of integration and measures of evolvability, namely conditioned covariance and evolutionary flexibility, were compared across anatomical groups within and among eight primate genera. As discussed in Section 2.4., these statistics provide estimates of the constraint, and thus potential of traits to respond to directional selection. We expect that there will be evidence that, especially for the basicranium, shoulder, and pelvis, relationships with each other will lead to constraints in the evolutionary potential of the traits within each anatomical region, following the results of Agosto and Auerbach (2021). Furthermore, it is expected that anatomical regions with direct functional anatomy and/or developmental relationships, such as the basicranium and shoulder girdle (see Matsuoka et al., 2005; Diogo and Wood, 2012), will have greater magnitudes of morphological integration than functionally unrelated traits within each taxon (see Cheverud, 1996; Wagner, 1996, Rolian, 2009). Among primates, we expect monkeys and apes to show a pattern of integration and evolvability where monkeys have greater integration and lower evolvability than apes for the same pairs of traits, following the broad patterns reported previously (Young, 2004; Rolian, 2009; Young et al., 2010; Lewton, 2012).

## 2. Materials and methods

### 2.1. Samples

Eight taxa are included in this study: seven catarrhines and one platyrrhine (Table 1). This sample represents a wide range of locomotor behaviors that vary in functional demands of the shoulder. When possible, each genus is represented by a single species. However, samples for both *Pongo* and *Papio* are represented by multiple species to achieve the minimum recommended sample sizes indicated by Grabowski and Porto’s (2017) ‘howmany’ R function (*n* ≥ 11 for 10 traits and *n* ≥ 21 for 20 traits for evolutionary flexibility; see Supplementary Online Materials [SOM] Table S1 for full results of the ‘howmany’ analysis). We combined species within each of these two genera based on results of a multivariate analysis of variance (MANOVA) of the traits between species, which showed no significant differences (*p* > 0.05) among traits used in this study among species of *Papio* or among species of *Pongo* (see SOM Table S2 for MANOVA results). Each taxon is represented by both males and females. Large numbers of missing teeth precluded the use of dental eruption to estimate age. Thus, we define adults as specimens with fusion of all long bone epiphyses. Only specimens with no visible pathologies affecting trait morphology were measured for this study. Additional information regarding the sample can be found in the SOM.

**Table 1.** Sample composition.

| Species | Males | Females | Total | Minimum Total<sup>a</sup> |
| --- | --- | --- | --- | --- |
| Platyrrhines |  |  |  |  |
| *Alouatta seniculus* | 18 | 23 | 41 | 24 |
| Catarrhines |  |  |  |  |
| Cercopithecoids |  |  |  |  |
| *Colobus guereza* | 27 | 26 | 53 | 33 |
| *Macaca mulatta* | 16 | 46 | 62 | 32 |
| *Papio* | 21 | 18 | 39 | 30 |
| *Pa. anubis* | 12 | 8 | - |  |
| *Pa. hamadryas* | 4 | 5 | - |  |
| *Pa. cynocephalus* | 5 | 5 | - |  |
| Hominoids |  |  |  |  |
| *Homo sapiens* | 56 | 60 | 116 | 105 |
| *Hylobates lar* | 28 | 24 | 52 | 42 |
| *Pan troglodytes* | 42 | 26 | 68 | 50 |
| *Pongo* | 21 | 31 | 52 | 32 |
| *P. abellii* | 2 | 7 | - |  |
| *P. pygmaeus* | 19 | 24 | - |  |

<sup>a</sup>These values account for individuals with missing data and represent the smallest possible sample size available for analyses.

### 2.2. Measurements

Twenty linear measurements (Fig. 1; SOM Table S3) were taken from the basicranium, shoulder girdle, pelvic girdle, and humerus. Morphological data were derived from the left side of each specimen. If the element or trait (in the case of the basicranium) from the left side was missing, damaged, or showed evidence of healed fracture, the right side of the specimen was measured for all elements. In 10 specimens, measurements were taken from both the right and left side of the specimen to comprise a ‘complete’ specimen due to missing elements/traits from both sides of the body.

**Figure 1.**
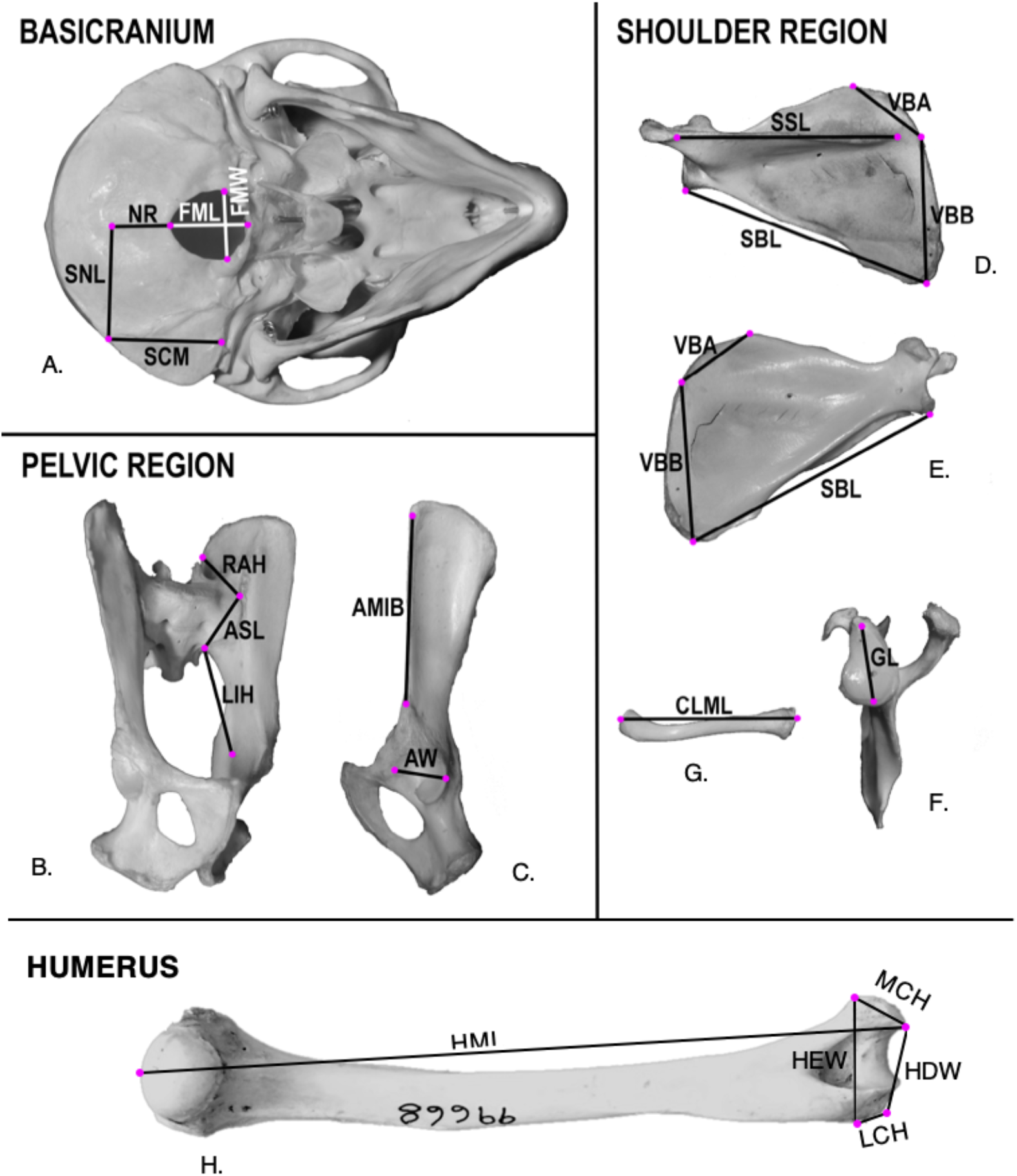
Linear dimensions collected from each anatomical region. A) caudal aspect of the basicranium; B) medial aspect of os coxa; C) lateral aspect of os coxa; D) dorsal aspect of scapula; E) ventral aspect of scapula; F) lateral aspect of scapula; G) cranial aspect of clavicle; H) ventral aspect of humerus. Measurements are demonstrated on a *Macaca mulatta* specimen and are homologous for all taxa. Trait descriptions and their developmental and functional correlates can be found in SOM Table S3.

We use the same selection of traits as Agosto and Auerbach (2021) for the basicranium, shoulder girdle, and pelvis. We also include additional traits from the distal humerus. Linear dimensions of the humerus are included in this study to characterize an element that has no established developmental or direct functional relationship with the basicranium or pelvis (e.g., no muscles in common and no direct association in movement), thus providing a more developmentally and functionally independent contrast with the results among the three other anatomical regions.

An Immersion G2X MiscroScribe digitizer was used to capture three-dimensional coordinates from the four anatomical regions (SOM Fig. S1; SOM Table S4). These coordinates were used to calculate interlandmark distances for the 20 dimensions used in this study (SOM Table S3). Three non-consecutive measurements were collected for each coordinate and linear measurement, and subjected to a repeated measures analysis of variance (RMANOVA) to assess intra-observer error. Results of the RMANOVA show no significant difference between repeated measures (*p* > 0.05), and <3% error for each element within taxa. All subsequent analyses were performed using the average of the three measurements.

### 2.3. Estimation of phenotypic variance-covariance matrices

Quantitative genetics analyses are performed using the additive genetic variances and covariances among traits (i.e., the G matrix; Lande, 1979; Hansen and Houle, 2008). However, these measures are notoriously difficult to obtain. For morphological traits, the phenotypic variances and covariances among traits have been shown to be close to proportional with their genetic counterparts (Roff, 1995, 1996; Cheverud, 1988; Roseman, 2012; Sodini et al., 2018). As such, the phenotypic variance-covariance (VCV) matrix, or P matrix, was substituted for the G matrix. VCV matrices were estimated using 20 traits across four anatomical regions (basicranium, shoulder girdle, pelvic girdle, and humerus) for each genus; the variance and covariance among traits for paired comparisons of regions were acquired from this matrix. Sources of variation not directly associated with the genotype-phenotype map (Wagner and Altenberg, 1996), in this case sex and species differences, were taken into account during matrix estimation by applying a MANOVA to the raw data and estimating the VCV matrices from the resulting residuals. Accounting for these sources of covariance is important because differences in the trait averages among groups can inflate the covariance among traits, therefore masking the true relationships among traits (Porto et al., 2013). While some studies have shown that subtle differences may exist in patterns of integration within and between elements intraspecifically between the sexes and with age (e.g., Mallard et al., 2017, Huseynov et al., 2017, Auerbach et al., 2018), we assume the differences between taxa are greater than intraspecific sexual dimorphism, which nonetheless is taken into account by using the residuals of the MANOVA.

All analyses of evolvability, conditioned covariance, flexibility, integration, and trait autonomy (see below) were performed using two approaches. In the first approach, the VCV matrices were standardized by the cross product of the trait means (Hansen and Houle, 2008); this method accounts for any mean-variance relationships that may be affecting the results, while maintaining size information within the VCV matrix. The second approach accounts for the influence of body size on the VCV matrices. Following Porto and colleagues (2013), size corrected VCV matrices (*R*) were obtained from the following relationship:

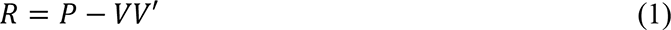

where *P* is the phenotypic VCV matrix, *V* is the size-related eigenvector and *V*′ is its transpose. Accounting for the effects of body size on the VCV matrices in this approach allows for the exploration of more subtle influences on trait variance and covariance, such as shared development and/or function, on the evolutionary potential of the anatomical regions included in this study.

Each VCV matrix (both mean standardized and size corrected) was also ‘noise corrected’ to minimize the effect small sample sizes and measurement error have on matrix estimation, especially in analyses that require matrix inversion (Twede and Hayden, 2004; Marroig et al., 2012). This correction minimizes the effect of small eigenvectors of the VCV matrix on our analyses; the small eigenvectors of a matrix often reflect random residual variance (Twede and Hayden, 2004; Houle et al., 2011), the effects of which are inflated due to matrix inversion in the calculation of conditioned covariance (see Equation 5) and trait autonomy (see Equation 6). Additional details on the noise correction can be found in SOM S1.

All analyses were performed in R v. 3.6.0 (R Core Team, 2019) using custom made scripts and the ‘evolvability’ package (Bolstad et al., 2014). Both data and code are available upon reasonable request.

### 2.4. Estimating evolvability, conditioned covariance, evolutionary flexibility, morphological integration, and trait autonomy

To analyze evolutionary potential and integration among sets of traits, we compared different pairings of traits (e.g., the shoulder girdle and pelvic girdle versus the shoulder girdle and basicranium). Each pairing consists of ten traits—five from each of the two regions—which is the minimum number of traits recommended by Grabowski and Porto (2017) for estimating measures of evolvability. Restricting the analysis to five traits from each region maximizes the sample (avoiding missing data) and avoids potential bias in estimates of evolvability and morphological integration that may be imposed due to unequal trait representation among anatomical regions.

To assess the consistency of the evolutionary covariance of the shoulder girdle with other anatomical regions, adaptations of Hansen and Houle’s (2008) evolvability equations and other measures of integration (Wagner, 1984, 1990) were used to assess two measures of morphological integration and evolvability both within and among taxonomic groups. The estimates of integration and evolvability for multivariate traits developed by Hansen and Houle (2008) were based on Lande (1979) and colleagues’ (1983) equation for estimating multivariate responses to directional selection

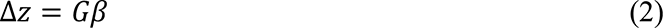

where Δ*z* is the average trait response to selection, *G* is the additive genetic VCV matrix (replaced by *P* in this study) and *β* is the directional selection gradient, which is a vector of partial regression coefficients of relative fitness on each trait in a multivariate system (Lande and Arnold, 1983, Hansen and Houle, 2008). The selection gradient essentially characterizes the direction and magnitude of selection in phenotypic space.

Hansen and Houle (2008) adapted this equation to estimate the capacity of a multivariate (i.e., multi-trait) population mean to respond to the directional selection. This measure, evolvability, provides an estimate of the potential for traits to evolve in phenotypic space, given their covariances with other traits and thus the potential constraints on patterns of variation characterized by the population’s VCV matrix (Rolian, 2009). Hansen and Houle (2008) define multivariate evolvability, *E*(*β*), as the length of the projection of evolutionary response vector (ι1z) on the selection gradient (*β*), divided by the norm of *β* (strength of directional selection):

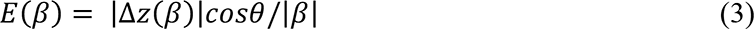

where cos θ is the angle between the response vector and the selection gradient, and the other variables are the same as in Equation 2. This equation provides a different measure of evolvability for each selection gradient, and therefore is unique in each direction of phenotypic space. The evolutionary potential of the *P* matrix can be assessed by computing its average evolvability over random selection gradients. In phenotypic space, the average trait additive variance is equal to the average eigenvalue (Hansen and Houle, 2008). As noted by Rolian (2009), evolutionary potential highly correlates with the total variance within a population, which prevents the comparison of average evolvabilities among species with unequal total variance. This study will use a related measure, evolutionary flexibility, to compare evolutionary potential across taxa.

Evolutionary flexibility, also referred to as the Directional Evolvability Index (Rolian, 2009), captures how closely the evolutionary response vector follows the selection gradient in phenotypic space (Marroig et al., 2009), and is measured as the cosine of the angle between these two vectors

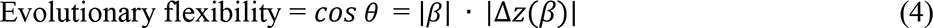

This is the portion of Hansen and Houle’s evolvability that measures the direction of the response of the population mean to the selection gradient and ignores the magnitude of evolutionary response, thus circumventing the issues with comparing evolvability between taxa with unequal total variance (Rolian, 2009). Essentially, evolutionary flexibility captures only the orientation of evolutionary response in phenotypic space (Marroig et al., 2009). Measures of evolutionary flexibility range from zero to one, where a measure of zero implies that the response vector is orthogonal to the direction of selection, and a measure of one is perfectly aligned with the selection vector (see Fig. 2). Thus, a value of one implies that traits are free to respond to any pressure of selection, while measures closer to zero indicate stronger constraint among traits.

**Figure 2.**
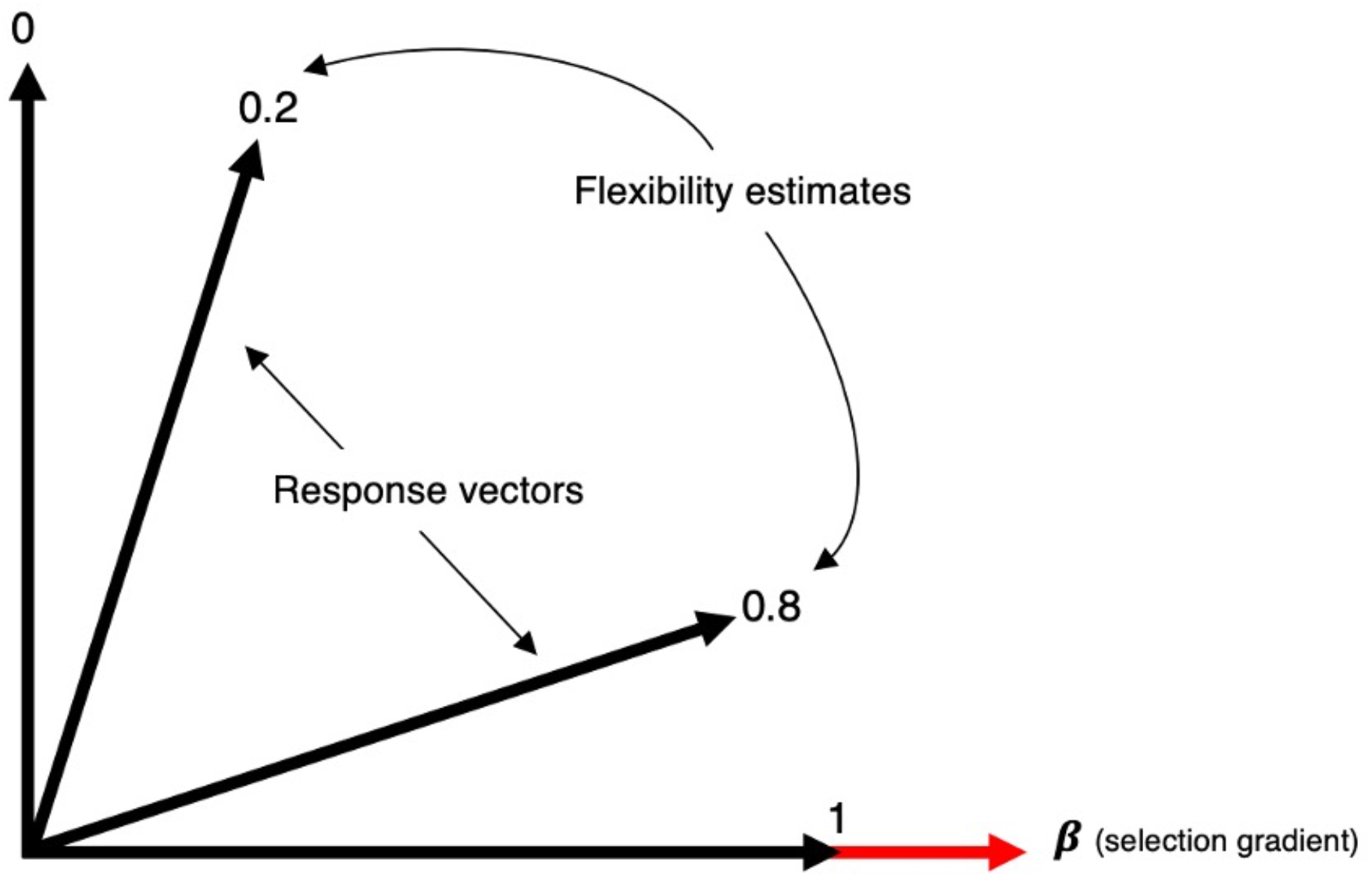
Comparison of response vectors (black) with different measures of evolutionary flexibility (numbers) against a selection gradient (*β*, red vector) in phenotypic space. Measures of evolutionary flexibility range from zero to one. A measure of one indicates that traits are free to respond in the direction of the selection gradient, and the response vector is parallel to the selection gradient. A measure of zero indicates high constraint among traits and they will be unable to respond in the direction of the selection gradient, and the response vector is orthogonal to the selection gradient.

The average evolutionary flexibility for each taxon was calculated using 1000 symmetrically distributed, randomly generated selection gradients. These were generated using the ‘evolvability’ package by independently selecting each element of the selection gradient from a zero-mean Gaussian distribution with a unit variance, and normalizing each selection gradient to a unit length by dividing by the norm (Bolstad et al., 2014). Thus, standardizing the length of the evolutionary response vector, which represents the magnitude of evolutionary change. The same set of 1000 selection gradients was used in all analyses. For each selection gradient, evolutionary flexibility was calculated as the cosine of the angle between the response vector (|Δ*z*(*β*)|) and the normalized selection gradient (|*β*|). The average evolutionary flexibility is the average of 1000 cosines. The 95% confidence interval (CI) for each analysis was calculated by bootstrapping the original data with replacement 1000 times, recomputing the evolutionary flexibility for each iteration of the bootstrapped sample (Manly, 1991), and calculating the confidence interval from the standard error of the total bootstrapped sample. Following Rolian (2009), statistical significance of the differences among taxa were determined by counting how many times the bootstrapped value in the CI of the smaller value exceeded that of the larger value and divided by the number of observations (1000). Statistically significant differences are based on α ≤ 0.05. Pairwise comparisons were performed between taxa.

Assessing the potential constraint among anatomical regions within taxa is achieved by estimating both the evolvability and the conditioned covariance among groups of traits. While evolvability provides an estimate of the ability of a group of traits to respond to directional selection, the conditioned covariance provides an estimate of which set of traits, if any, impose greater genetic constraint over others. Measures of conditioned covariance estimate the potential constraint a group of traits under stabilizing selection (x) imposes on the ability of a set of traits (y) to respond to directional selection (Hansen and Houle, 2008). This provides a general measure of the genetic constraint among traits by estimating the conditional genetic covariance of y given x (Hansen and Houle, 2008). Conditioned covariance is calculated using the following equation:

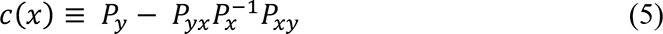

where *P_y_* is the phenotypic variance among a set of traits (y), *P_x_* is the phenotypic variance of a set of traits under stabilizing selection (x), and *P_xy_* and *P_yx_* are the row and column vectors of covariances between x and y (Hansen, 2003; Hansen et al., 2003; Hansen and Houle, 2008). We calculate the conditioned covariance for each pairing of anatomical regions, with each region standing in for both x and y. Confidence intervals were calculated using the bootstrapping technique discussed above.

Morphological integration among anatomical regions for each taxon was assessed using two methods: the variance of eigenvalues (VE) and trait autonomy (TA). VE provides a measure of the strength of the covariance among traits within a population (Rolian, 2009). Higher measures of VE indicate that a small number of principal components with large eigenvalues explain most of the variance in the data. Stronger integration among traits suggests greater covariance among them. The VE method was developed for correlation matrices (Wagner, 1984), and Rolian (2009) demonstrated its application to VCV matrices and how it can be adjusted so measures of VE are proportional across taxa of unequal variances; thus, allowing their comparison both among anatomical regions within a taxon, as well as between taxa. This adjustment entails standardizing eigenvalues by the trace of the VCV matrix, so that each eigenvalue is expressed as a portion of the total variance, where the sum of all eigenvalues is one for each taxon. Confidence intervals and statistical differences among taxa were calculated using the same bootstrap strategy described for evolutionary flexibility (1000 iterations).

TA reflects the magnitude of morphological integration by estimating the amount of genetic or phenotypic variance of a trait that is independent of the traits with which it covaries (Hansen and Houle, 2008). This measure therefore quantifies the constraint on a trait with respect to a set of traits. Trait autonomy for a single trait z_j_, with respect to all others, is given by:

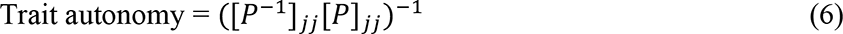

where *P^-1^* is the inverse of the *P* matrix, and _jj_ distinguishes the jth trait along the diagonal of the matrix (Hansen and Houle, 2008). Estimates of TA range from zero to one, and can be conceptualized as the percentage of phenotypic variance that is free to respond to selection without the influence of other traits; in essence, this measure represents the size of the effect that shared covariances have on the potential for a trait to independently respond to directional selection. And so, the average autonomy across a suite of traits provides a measure of how parcellated the data is (Wagner and Altenberg, 1996). Traits with a lower autonomy have a greater proportion of their potential variance ‘locked up’ in their covariance with other traits (Rolian, 2009). Like evolvability, estimates of TA cannot be compared between taxa of unequal variance, thus we only compare values of TA between anatomical regions within a taxon and discuss the patterns of TA between taxa. Confidence intervals and significance among comparisons of trait autonomy use the same resampling strategy mentioned above (1000 iterations).

## 3. Results

### 3.1. Evolvability and conditioned covariance

Within each genus, estimates of evolvability between anatomical regions are all similar (see SOM Tables S5 and S6). Due to this trend, comparisons between anatomical regions within each genus were based on the estimates of conditioned covariance. This measure provides a more nuanced look at the constraint between sets of traits compared to evolvability, which estimates the potential evolutionary constraint among a group of traits as a whole. Estimates of conditioned covariance, standard errors, and 95% confidence intervals calculated with size corrected VCV matrices are presented in Table 2 and in SOM Figure S2. Results based on mean standardized VCV matrices are presented in SOM Table S7. Much like evolvability, point estimates of conditioned covariance cannot be compared between groups with unequal variance (which is inherent in our sample), and so we instead compare between the genera the patterns of conditioned covariance observed within genera as opposed to the actual values of the point estimates themselves.

**Table 2.**
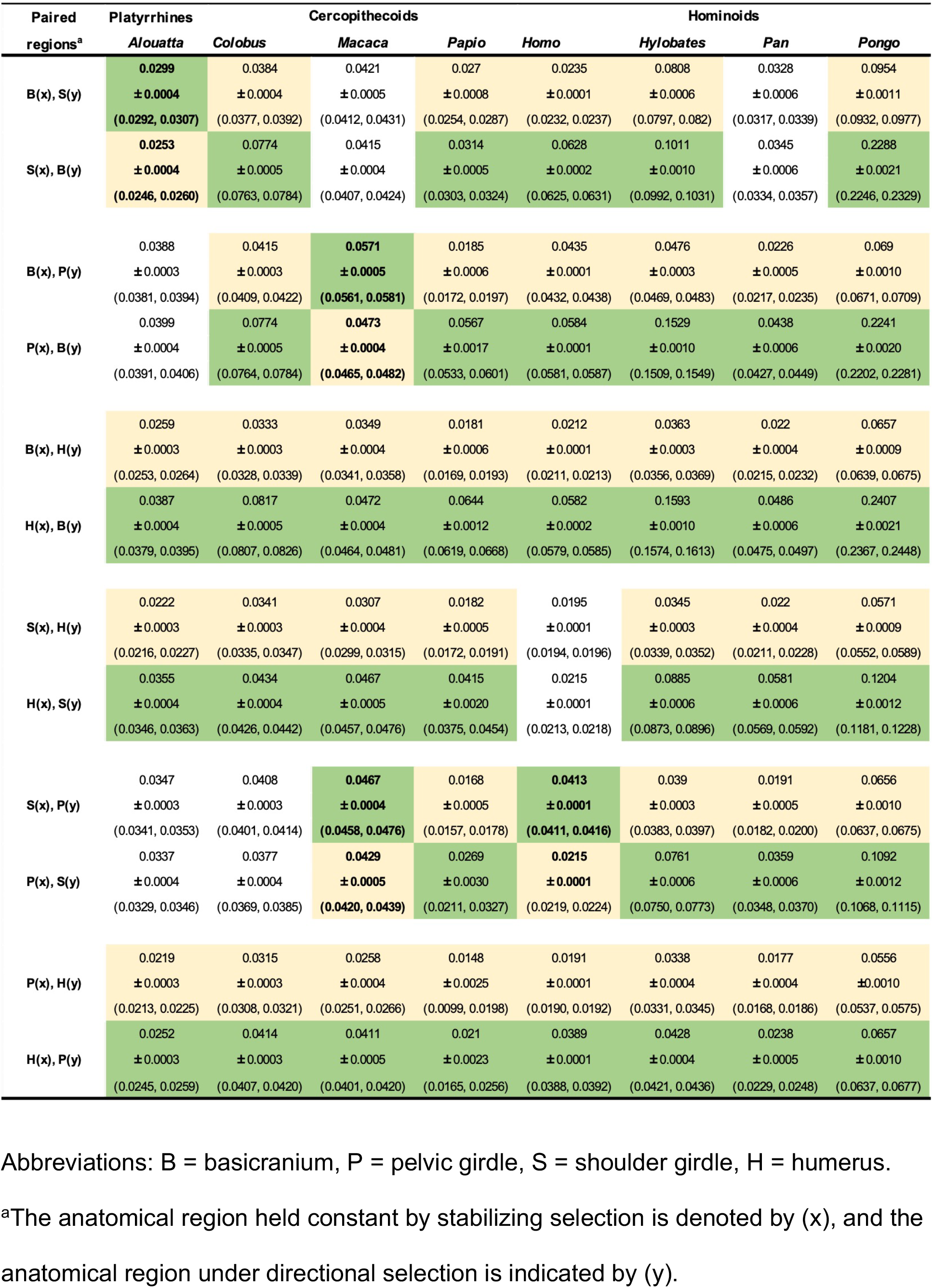
Mean conditioned covariance, standard error (indicated by ±), and 95% confidence intervals (in parentheses) for paired anatomical regions for each genus using size corrected variance covariance matrices. Among comparisons of anatomical regions, cells colored light yellow indicate the lower conditioned covariance, whereas cells colored green have a higher conditioned covariance. Uncolored cells indicate comparisons where the conditioned covariances are similar. Lower values reflect smaller responses to directional selection, and thus more constraint being imposed by the set of traits under stabilizing selection. Analyses that deviate from the primary pattern among genera have bolded values.

Both the mean standardized (SOM Table S7) and size corrected (Table 2; SOM Figure S2) results show a fairly consistent pattern among taxa with some exceptions; both the patterns and taxa that deviate from them are consistent between both sets of results. In most taxa, traits of the basicranium have a potential to impose greater constraint on all other anatomical regions assessed than those regions can impose on the basicranium. Exceptions are observed in *Macaca*, *Alouatta*, and *Pan*. The shoulder girdle has the potential to impose greater constraint on the pelvis and humerus than these anatomical regions may impose on it. Deviations from this trend are observed in *Homo*, *Macaca*, *Alouatta*, and *Colobus*. All anatomical regions have the potential to impose more constraint on the humerus than the humerus can impose on those regions.

### 3.2. Evolutionary flexibility

As noted in Section 2.4., unlike conditioned covariance, estimates of evolutionary flexibility may be compared between taxa for the same traits, as it focuses on the direction and not the magnitude of the evolutionary response vector. The mean evolutionary flexibility among paired anatomical regions, as well as the 95% confidence intervals and standard errors, for the VCV matrices accounting for size are presented in Table 3. Results for evolutionary flexibility estimates derived from the mean standardized VCV matrices (see SOM Table S8) show an overall inverse trend associated with body size; smaller genera (typically monkeys) have larger estimates of flexibility, and thus greater potential to respond to directional selection, compared to larger genera (apes). Body size has been shown to be an important factor in mammalian evolution (Marroig et al., 2009; Porto et al., 2013), one that can overwhelm the signals presented by local factors that may also affect trait evolution, such as those of interest to this study (shared development and function among anatomical regions). As the results derived from the mean standardized VCV matrices appear to be motivated by body size, we have reported these in the SOM (Tables S8, S11, S12) and hereafter, all results and discussion are based on the VCV matrices that take size into account. We will note that these results represent the portion of evolutionary potential that is due to factors other than those related to body size (though some may relate to body size), and not the total evolutionary potential of these regions (which would include body size).

**Table 3.**
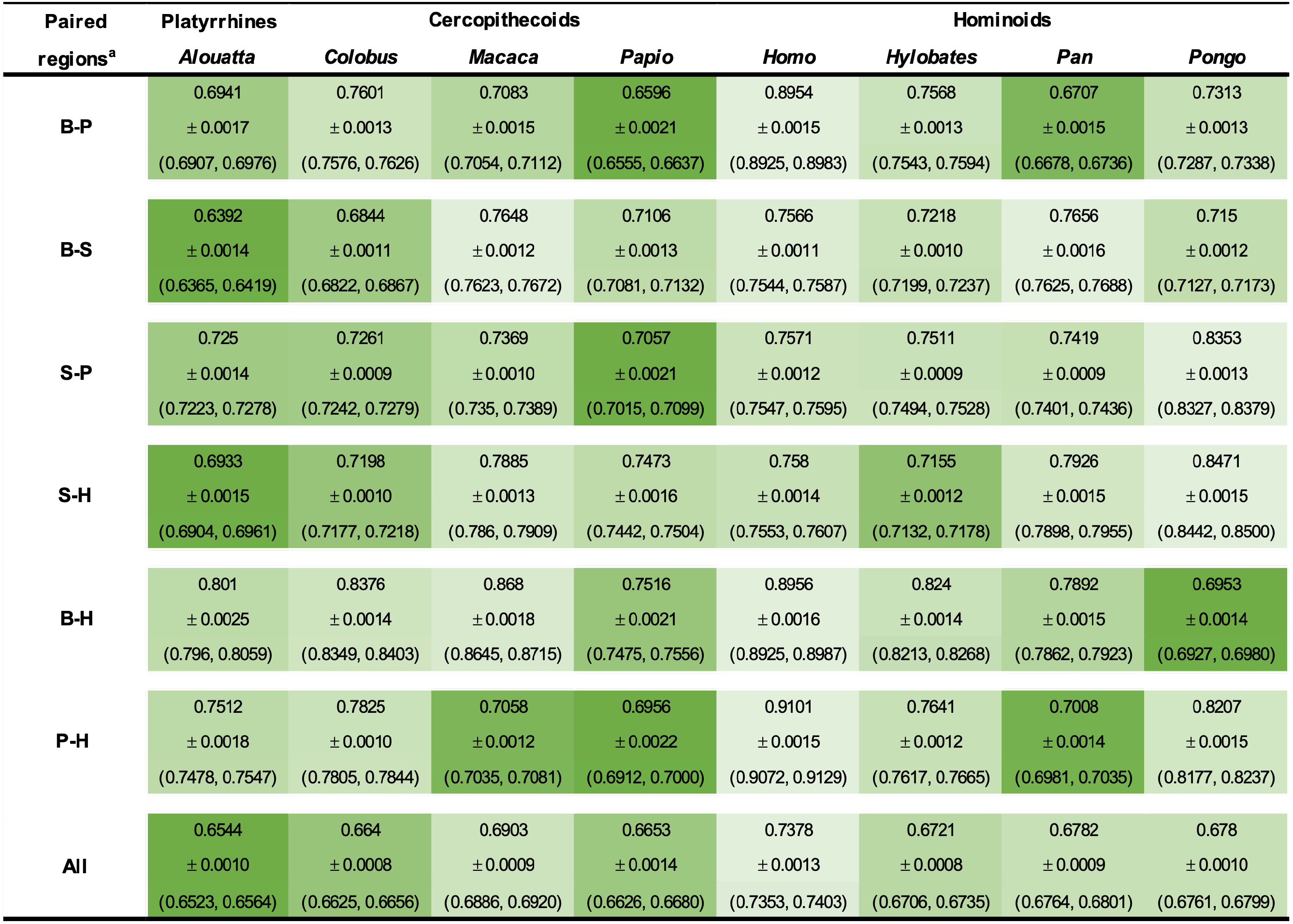
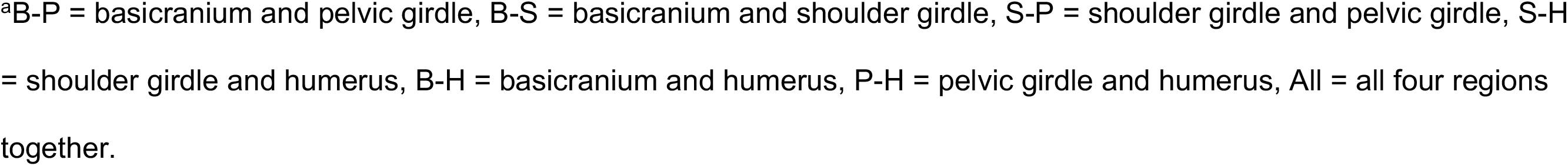
Mean evolutionary flexibility, standard error (indicated by ±), and 95% confidence intervals (in parentheses) for paired anatomical regions among genera using size corrected variance covariance matrices. Higher values reflect greater flexibility, and thus more evolvability for that grouping of traits compared among genera (rows). Colors indicate the level of evolutionary flexibility compared between genera for each paired anatomical region: lighter shades of green indicate less constraint and darker shades of green indicate more constraint among regions.

| Paired<br>regions<sup>a</sup> | Platyrrhines | Cercopithecoids |  |  | Hominoids |  |  |  |
| --- | --- | --- | --- | --- | --- | --- | --- | --- |
|  | *Alouatta* | *Colobus* | *Macaca* | *Papio* | *Homo* | *Hylobates* | *Pan* | *Pongo* |
| **B-P** | 0.6941 | 0.7601 | 0.7083 | 0.6596 | 0.8954 | 0.7568 | 0.6707 | 0.7313 |
|  | ± 0.0017 | ± 0.0013 | ± 0.0015 | ± 0.0021 | ± 0.0015 | ± 0.0013 | ± 0.0015 | ± 0.0013 |
|  | (0.6907, 0.6976) | (0.7576, 0.7626) | (0.7054, 0.7112) | (0.6555, 0.6637) | (0.8925, 0.8983) | (0.7543, 0.7594) | (0.6678, 0.6736) | (0.7287, 0.7338) |
| **B-S** | 0.6392 | 0.6844 | 0.7648 | 0.7106 | 0.7566 | 0.7218 | 0.7656 | 0.715 |
|  | ± 0.0014 | ± 0.0011 | ± 0.0012 | ± 0.0013 | ± 0.0011 | ± 0.0010 | ± 0.0016 | ± 0.0012 |
|  | (0.6365, 0.6419) | (0.6822, 0.6867) | (0.7623, 0.7672) | (0.7081, 0.7132) | (0.7544, 0.7587) | (0.7199, 0.7237) | (0.7625, 0.7688) | (0.7127, 0.7173) |
| **S-P** | 0.725 | 0.7261 | 0.7369 | 0.7057 | 0.7571 | 0.7511 | 0.7419 | 0.8353 |
|  | ± 0.0014 | ± 0.0009 | ± 0.0010 | ± 0.0021 | ± 0.0012 | ± 0.0009 | ± 0.0009 | ± 0.0013 |
|  | (0.7223, 0.7278) | (0.7242, 0.7279) | (0.735, 0.7389) | (0.7015, 0.7099) | (0.7547, 0.7595) | (0.7494, 0.7528) | (0.7401, 0.7436) | (0.8327, 0.8379) |
| **S-H** | 0.6933 | 0.7198 | 0.7885 | 0.7473 | 0.758 | 0.7155 | 0.7926 | 0.8471 |
|  | ± 0.0015 | ± 0.0010 | ± 0.0013 | ± 0.0016 | ± 0.0014 | ± 0.0012 | ± 0.0015 | ± 0.0015 |
|  | (0.6904, 0.6961) | (0.7177, 0.7218) | (0.786, 0.7909) | (0.7442, 0.7504) | (0.7553, 0.7607) | (0.7132, 0.7178) | (0.7898, 0.7955) | (0.8442, 0.8500) |
| **B-H** | 0.801 | 0.8376 | 0.868 | 0.7516 | 0.8956 | 0.824 | 0.7892 | 0.6953 |
|  | ± 0.0025 | ± 0.0014 | ± 0.0018 | ± 0.0021 | ± 0.0016 | ± 0.0014 | ± 0.0015 | ± 0.0014 |
|  | (0.796, 0.8059) | (0.8349, 0.8403) | (0.8645, 0.8715) | (0.7475, 0.7556) | (0.8925, 0.8987) | (0.8213, 0.8268) | (0.7862, 0.7923) | (0.6927, 0.6980) |
| **P-H** | 0.7512 | 0.7825 | 0.7058 | 0.6956 | 0.9101 | 0.7641 | 0.7008 | 0.8207 |
|  | ± 0.0018 | ± 0.0010 | ± 0.0012 | ± 0.0022 | ± 0.0015 | ± 0.0012 | ± 0.0014 | ± 0.0015 |
|  | (0.7478, 0.7547) | (0.7805, 0.7844) | (0.7035, 0.7081) | (0.6912, 0.7000) | (0.9072, 0.9129) | (0.7617, 0.7665) | (0.6981, 0.7035) | (0.8177, 0.8237) |
| **All** | 0.6544 | 0.664 | 0.6903 | 0.6653 | 0.7378 | 0.6721 | 0.6782 | 0.678 |
|  | ± 0.0010 | ± 0.0008 | ± 0.0009 | ± 0.0014 | ± 0.0013 | ± 0.0008 | ± 0.0009 | ± 0.0010 |
|  | (0.6523, 0.6564) | (0.6625, 0.6656) | (0.6886, 0.6920) | (0.6626, 0.6680) | (0.7353, 0.7403) | (0.6706, 0.6735) | (0.6764, 0.6801) | (0.6761, 0.6799) |

<sup>a</sup>B-P = basicranium and pelvic girdle, B-S = basicranium and shoulder girdle, S-P = shoulder girdle and pelvic girdle, S-H = shoulder girdle and humerus, B-H = basicranium and humerus, P-H = pelvic girdle and humerus, All = all four regions together.

Broadly, apes consistently present the highest mean evolutionary flexibility values for all combinations of traits compared to the two groups of monkeys. Traits are collectively least constrained in *Homo*, especially among pairings that lack direct functional and developmental relationships: basicranium, pelvic girdle, and humerus. Conversely, pairings among these three anatomical regions are more constrained in *Pan*, where the mean evolutionary flexibility estimates are more similar to those observed in monkeys. Among the monkeys, mean evolutionary flexibility estimates are collectively highest in *Macaca*, and lowest in *Papio*. Different from all other taxa, *Pongo* shows the greatest potential constraint between the basicranium and humerus. Significance between mean estimates of evolutionary flexibility among taxa are presented in the SOM Table S9. Notably, significant differences in the estimates of evolutionary flexibility are most frequently observed between *Homo* and other taxa.

### 3.3. Morphological integration

The index of integration, given by the trace-adjusted eigenvalues of the size corrected VCV matrix, standard errors, and the 95% confidence intervals for each paired grouping of anatomical regions, can be found in Table 4 for the VE and in SOM Table S10 for TA; mean standardized results are presented in SOM Table S11 for VE and SOM Table S12 for TA. For measures of VE, larger values indicate greater integration and therefore greater potential constraint among traits, whereas smaller values of trait autonomy indicate less independence, and therefore greater potential evolutionary constraint, among traits. Statistical significance between pairs of anatomical regions among taxa can be found in SOM Table S9.

**Table 4.**
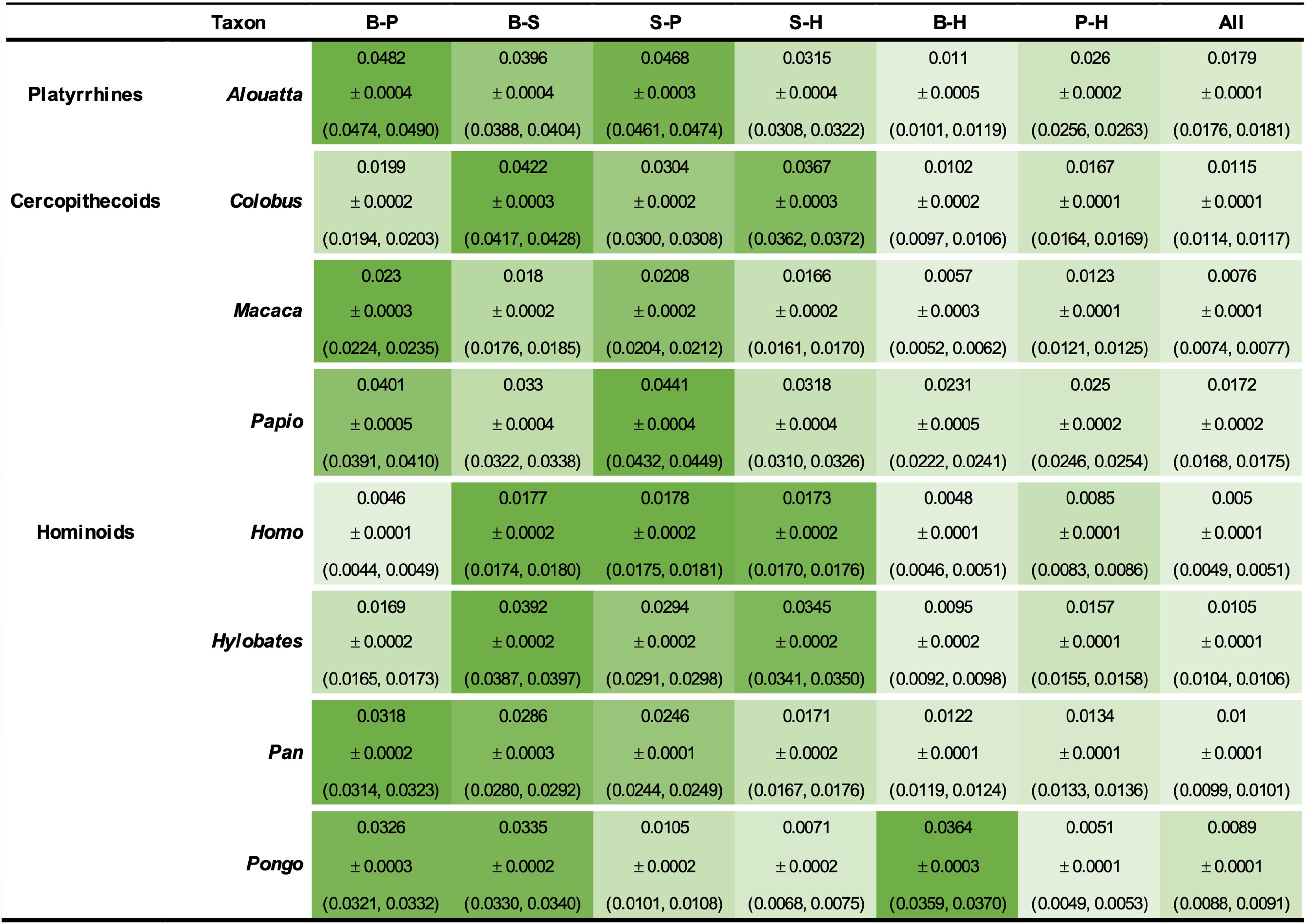
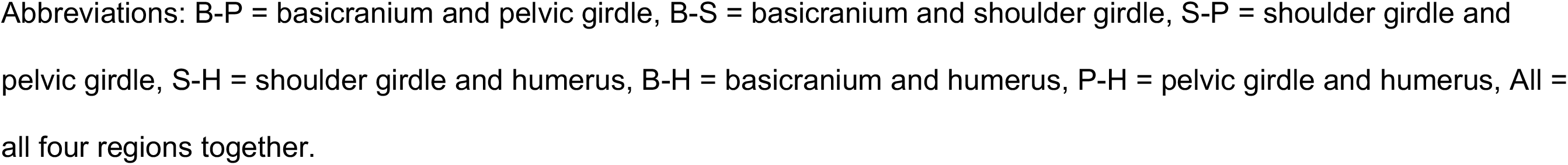
Mean variance of the eigenvalues (VE), standard errors (indicated by ±), and 95% confidence intervals (in parentheses) by taxa for pairwise trait groupings among anatomical regions (columns) using size corrected variance covariance matrices. Greater VE values reflect stronger integration among the traits. Comparisons among traits within genera are in rows. Colors reflect the degree of integration, and thus constraint, between pairs of anatomical regions within each taxon: lighter shades of green indicate weaker integration among traits and darker shades of green indicate stronger integration among traits.

| Taxon |  | B-P | B-S | S-P | S-H | B-H | P-H | All |
| --- | --- | --- | --- | --- | --- | --- | --- | --- |
| Platyrrhines | *Alouatta* | 0.0482 | 0.0396 | 0.0468 | 0.0315 | 0.011 | 0.026 | 0.0179 |
|  |  | ± 0.0004 | ± 0.0004 | ± 0.0003 | ± 0.0004 | ± 0.0005 | ± 0.0002 | ± 0.0001 |
|  |  | (0.0474, 0.0490) | (0.0388, 0.0404) | (0.0461, 0.0474) | (0.0308, 0.0322) | (0.0101, 0.0119) | (0.0256, 0.0263) | (0.0176, 0.0181) |
| Cercopithecoids | *Colobus* | 0.0199 | 0.0422 | 0.0304 | 0.0367 | 0.0102 | 0.0167 | 0.0115 |
|  |  | ± 0.0002 | ± 0.0003 | ± 0.0002 | ± 0.0003 | ± 0.0002 | ± 0.0001 | ± 0.0001 |
|  |  | (0.0194, 0.0203) | (0.0417, 0.0428) | (0.0300, 0.0308) | (0.0362, 0.0372) | (0.0097, 0.0106) | (0.0164, 0.0169) | (0.0114, 0.0117) |
|  | *Macaca* | 0.023 | 0.018 | 0.0208 | 0.0166 | 0.0057 | 0.0123 | 0.0076 |
|  |  | ± 0.0003 | ± 0.0002 | ± 0.0002 | ± 0.0002 | ± 0.0003 | ± 0.0001 | ± 0.0001 |
|  |  | (0.0224, 0.0235) | (0.0176, 0.0185) | (0.0204, 0.0212) | (0.0161, 0.0170) | (0.0052, 0.0062) | (0.0121, 0.0125) | (0.0074, 0.0077) |
|  | *Papio* | 0.0401 | 0.033 | 0.0441 | 0.0318 | 0.0231 | 0.025 | 0.0172 |
|  |  | ± 0.0005 | ± 0.0004 | ± 0.0004 | ± 0.0004 | ± 0.0005 | ± 0.0002 | ± 0.0002 |
|  |  | (0.0391, 0.0410) | (0.0322, 0.0338) | (0.0432, 0.0449) | (0.0310, 0.0326) | (0.0222, 0.0241) | (0.0246, 0.0254) | (0.0168, 0.0175) |
| Hominoids | *Homo* | 0.0046 | 0.0177 | 0.0178 | 0.0173 | 0.0048 | 0.0085 | 0.005 |
|  |  | ± 0.0001 | ± 0.0002 | ± 0.0002 | ± 0.0002 | ± 0.0001 | ± 0.0001 | ± 0.0001 |
|  |  | (0.0044, 0.0049) | (0.0174, 0.0180) | (0.0175, 0.0181) | (0.0170, 0.0176) | (0.0046, 0.0051) | (0.0083, 0.0086) | (0.0049, 0.0051) |
|  | *Hylobates* | 0.0169 | 0.0392 | 0.0294 | 0.0345 | 0.0095 | 0.0157 | 0.0105 |
|  |  | ± 0.0002 | ± 0.0002 | ± 0.0002 | ± 0.0002 | ± 0.0002 | ± 0.0001 | ± 0.0001 |
|  |  | (0.0165, 0.0173) | (0.0387, 0.0397) | (0.0291, 0.0298) | (0.0341, 0.0350) | (0.0092, 0.0098) | (0.0155, 0.0158) | (0.0104, 0.0106) |
|  | *Pan* | 0.0318 | 0.0286 | 0.0246 | 0.0171 | 0.0122 | 0.0134 | 0.01 |
|  |  | ± 0.0002 | ± 0.0003 | ± 0.0001 | ± 0.0002 | ± 0.0001 | ± 0.0001 | ± 0.0001 |
|  |  | (0.0314, 0.0323) | (0.0280, 0.0292) | (0.0244, 0.0249) | (0.0167, 0.0176) | (0.0119, 0.0124) | (0.0133, 0.0136) | (0.0099, 0.0101) |
|  | *Pongo* | 0.0326 | 0.0335 | 0.0105 | 0.0071 | 0.0364 | 0.0051 | 0.0089 |
|  |  | ± 0.0003 | ± 0.0002 | ± 0.0002 | ± 0.0002 | ± 0.0003 | ± 0.0001 | ± 0.0001 |
|  |  | (0.0321, 0.0332) | (0.0330, 0.0340) | (0.0101, 0.0108) | (0.0068, 0.0075) | (0.0359, 0.0370) | (0.0049, 0.0053) | (0.0088, 0.0091) |

Abbreviations: B-P = basicranium and pelvic girdle, B-S = basicranium and shoulder girdle, S-P = shoulder girdle and pelvic girdle, S-H = shoulder girdle and humerus, B-H = basicranium and humerus, P-H = pelvic girdle and humerus, All = all four regions together.

Comparisons of morphological integration between pairs of anatomical regions within genera follow a fairly consistent pattern for both measures of VE and TA. As such, we focus our discussion of morphological integration on the size corrected VE results in this section. *Homo* has the lowest morphological integration when traits from all four anatomical regions are considered together, whereas *Alouatta* has the highest values of VE. Overall, apes tend to be less integrated than monkeys. One notable exception is *Macaca*, which has the second lowest global measures of morphological integration among taxa in this study. Morphological integration is lowest between the basicranium and humerus in all taxa except *Pongo*, where, paralleling the flexibility estimates, it is highest. For most taxa, the highest morphological integration, and thus lowest genetic independence, is observed when traits of the shoulder girdle are paired with traits from the basicranium or pelvic girdle.

## 4. Discussion

### 4.1. Evolutionary constraint among anatomical regions

The results present remarkably consistent patterns among primates of both morphological integration and conditioned covariance in paired groups of traits from the basicranium, shoulder girdle, pelvic girdle, and humerus. These results support our prediction that evolutionary constraint among the shoulder girdle, pelvic girdle, and basicranium is present among all primate taxa included in this study. Our results further uphold previous examinations of evolutionary constraint among the primate shoulder girdle, basicranium, and pelvic girdle (see Agosto and Auerbach, 2021). Moreover, the patterns in our results collectively demonstrate a reduction of constraint between anatomical regions that do not have direct functional or developmental relationships, such as the basicranium and humerus. A broad conclusion is that the primate shoulder region, as a whole, is consistently among the least autonomous and more morphologically integrated anatomical regions considered in this study.

Evolutionary constraint among traits will, in part, be due to the strength of the underlying covariances (genetic, functional, and developmental) among them (Cheverud, 1996; Hansen and Houle, 2008; Rolian, 2009, 2014). Evaluating isolated traits or skeletal elements within an evolutionary context would fail to account for these relationships among traits, and implicitly models the evolution of those traits as if they were not covariant with other aspects of anatomy. Previous studies of the primate shoulder girdle have assessed this region and its traits individually (e.g. Ashton et al., 1965; Trinkaus et al., 2014; Selby and Lovejoy, 2017), and thus, have worked under the assumption of its independent evolution. However, independent evolution of the traits of the shoulder girdle is not supported by the findings of Agosto and Auerbach (2021), nor by the results of this study. Among our sample, we observe a consistent pattern of a cascade-like effect of constraint from superior to inferior in the axial skeleton, with the traits of the axial skeleton imposing more constraint on appendicular traits than they impose on it. This pattern may reflect differences in developmental timing or a myriad of other overlapping factors beyond the scope of our investigation, and that are likely difficult to model (see Hallgrímsson et al., 2009, and Hallgrímsson et al., 2019), but nonetheless warrant further investigation.

While an explanation for this pattern of constraint among anatomical regions is unclear, the observed patterns of conditioned covariance in this study demonstrate evolutionary non-independence among the shoulder girdle, basicranium, pelvis, and humerus. The patterns of conditioned covariance only show directionality of evolutionary constraint between paired regions, whereas the patterns of integration and autonomy demonstrate differences in the potential evolutionary constraint across different pairings of anatomical regions. Within genera, the pattern of higher integration and lower autonomy observed among the pairings of anatomical regions involving the shoulder girdle indicates this region displays the most potential evolutionary constraint with other regions. Thus, this further supports evolutionary non-independence of the shoulder girdle in primates.

Shared trait responses to directional selection are ultimately influenced by the developmental and functional relationships among them. This occurs through increased pleiotropy on a population level among traits that are more strongly integrated due to shared developmental processes or engagement in similar functional demands within individuals (Cheverud, 1996; Wagner, 1996). The relationship among shared function, the strength of integration, and the constraints these may have on trait response to directional selection have been empirically demonstrated among the autopods (Rolian, 2009) and limbs (Young et al., 2010) of primates. These studies show that traits from anatomical regions that engage in similar functional demands are more strongly integrated, and overall, less evolvable than functionally independent traits. The patterns of morphological integration among pairs of anatomical regions observed in this study bear out these expectations.

There is evidence that the shoulder and basicranium share developmental pathways (Matsuoka et al., 2005; see also Lieberman et al., 2000, and Valasek et al., 2010). In many primate species, these two regions are physically connected by muscles of the neck and shoulder, or at least by ligaments and connective tissues (Richmond et al., 2001; Diogo and Wood, 2012). In addition, the shoulder and pelvic girdle are developmentally linked as analogous morphological regions (limb girdles) (Sears et al., 2015), and in quadrupedal species these share some broad locomotor functions. With both developmental and functional sources of covariance, it is unsurprising that the strongest relationships are often found among these regions. As noted above, in most taxa, integration between the distal humerus and these other three regions is lower than those amongst those three regions. This may in part be an artifact of the use of distal articular dimensions for the humerus; the proximal end of the humerus provides attachment sites for several muscles originating from the scapula (Diogo and Wood, 2012), and was excluded because it was thought to be redundant with the traits selected from the scapula for the shoulder girdle. It is worth noting that the distal humerus shows high covariance with body size among primates (Ruff, 2003), and its lower integration and less potential effect on the evolution of other traits, as indicated by our findings, argues against body size as being a primary motivator behind the results we report.

Compared to other pairings of anatomical regions, the basicranium and humerus have the lowest morphological integration, and thus autonomy, among their traits within each taxon. This is expected given the lack of known direct functional or developmental relationships between traits of the basicranium and distal humerus. However, we would be remiss to not highlight one notable exception: *Pongo* is the only genus that substantially deviates from this broad primate pattern of integration, though orangutans are similar to other apes in being less integrated than monkeys.

Within *Pongo*, the basicranium and humerus are among the most integrated anatomical regions, along with the basicranium and pelvic girdle, and therefore would experience more constraint among these regions than the other pairings of anatomical regions. More surprisingly, the shoulder region and humerus are among the least constrained pairings of anatomical regions, whereas these regions are more integrated in all other genera. This may suggest that *Pongo* ‘broke’, or relaxed, the evolutionary and developmental mechanism underlying the high covariance between these regions observed in other primate genera. Among apes, orangutans have an exceptionally high intermembral index for their size due to disproportionately long humeri (Jungers, 1985). Forelimb length in *Pongo* scales positively with body size, and to a greater degree than what is observed among other primates (Jungers, 1985). It is possible the relationship among the basicranium and humerus observed here is an artifact of the size correction in the VCV matrix. Most likely, however, the relationship we are observing between these seemingly unrelated anatomical regions may be due to unrelated similarities in the covariance structures of the basicranium and humerus that results in their mutual constraint. The exact cause of this relationship is unknown and warrants further investigation with measurements from other skeletal regions, as well as a better understanding of orangutan development compared with other apes.

### 4.2. Function and evolutionary flexibility

The second question asked in this study—examining functional variation in the use of the upper limb and evidence for corresponding amounts of evolutionary constraint among primates—concerns the results of the evolutionary flexibility of paired anatomical regions among primate taxa. Recall that this measure estimates evolutionary potential as the orientation of the response vector in relation to the directional selection vector, and not the magnitude of evolutionary change among traits. While estimates of integration and conditioned covariance suggest similar patterns of constraint across paired anatomical regions, the pattern of evolutionary flexibility within these pairings vary among primate taxa. Furthermore, these patterns do not strictly correspond to functional differences of the upper limb in primates and bear further examination.

The sample used in this study comprises a group of primate genera that represent a range of different functional demands placed on the shoulder. Given the relationship between shoulder girdle morphology and functional demands (Inman et al., 1944; Ashton and Oxnard, 1964a; Ashton et al., 1965; Oxnard, 1967; Ashton et al., 1976; Roberts, 1974; Larson, 1993, 1995, 2015; Irwin and Larson 2000), we had anticipated that the pattern of evolutionary flexibility among primates would correspond to the functional variation observed among taxa. However, this expectation is not borne out. Evolutionary flexibility among the gestalt of traits from the four anatomical regions are quite similar among all taxa, save *Homo*, which has the highest estimate of evolutionary flexibility, generally significantly greater than all other primates (see SOM Table S9), and thus greatest overall evolutionary potential of primates sampled. Comparing evolutionary flexibility estimates across genera for paired anatomical regions also does not support a function-driven explanation for the observed patterns, as different evolutionary potential is demonstrated for genera that engage in similar locomotor strategies. *Papio* and *Macaca* are considered ‘true’ terrestrial quadrupeds (see Meyers et al., 2019), and yet the estimates of evolutionary flexibility do not consistently cluster together within either paired anatomical regions or the gestalt of all traits (Table 3; SOM Table S9). For example, *Papio* exhibits a degree of evolutionary potential that is more similar with *Alouatta*, an arboreal quadruped, and *Pongo*, a large-bodied suspensory primate, than with *Macaca* for the shoulder girdle and pelvis, and basicranium and humerus pairings, respectively. We also do not observe a consistent pattern among the arboreal quadrupedal primates, *Alouatta* and *Colobus*, or among the suspensory primates, *Hylobates* and *Pongo*.

Although the pattern of evolutionary potential does not follow functional variation in the upper limb, or wide locomotor patterns among genera, it does appear to follow a broader evolutionary stroke separating monkeys and apes. Apes are generally less constrained and have greater evolutionary potential than monkeys, as borne out in both the measures of integration and constraint, as well as evolutionary flexibility in our results. Reduced constraint among apes, and especially humans, has previously been demonstrated in the limbs (Young et al., 2010) and scapula (Young, 2004). Our results demonstrate support that the evolutionary shift that occurred in the primate bauplan during the separation of monkeys and apes resulted in the breaking of the pattern of constraints among traits and anatomical regions that had previously been co-constrained in primates.

The observed variation in evolutionary flexibility estimates among primate genera is noteworthy, especially in light of the generally consistent results found among primates in the morphological integration and trait autonomy estimates. The most notable of these is observed in *Pan*. Concerning morphological integration and trait autonomy, *Pan* resembles the other hominoids. However, the patterns of evolutionary flexibility differ markedly from the other hominoid genera. This is especially surprising given the close phylogenetic relationship between *Pan* and *Homo*. While apes generally experience relaxed constraint among anatomical regions compared to monkeys, *Pan* breaks this pattern in the pairings between the basicranium and humerus, the basicranium and pelvis, and the pelvis and humerus. Among these three pairs of anatomical regions, *Pan* is among the most constrained genus that we assessed and has more monkey-like estimates of evolutionary flexibility. Thus, the pattern of evolutionary potential for *Pan* appears to represent a mix of both ape and monkey trends. This may be due to a combination of anatomical regions being constrained by shared history among hominoids, and the changes in co-constraint among traits during the separation of monkeys and apes allowing for the exploration of new evolutionary compromises in *Pan*. This result may also reflect the unique form of vertical climbing and knucklewalking *Pan* employs compared to other hominoids. Chimpanzees do not represent the only oddity among primates, as estimates of evolutionary flexibility that do not adhere to the monkey or ape trend are also observed in *Pongo*, *Colobus*, and *Macaca*. Altogether, the combined results in the patterns of morphological integration and evolutionary flexibility suggest we cannot make assumptions of how locomotion affects the evolution of traits under a multi-trait model of responses to selection.

One important caveat to our results is the analyses herein cannot show whether the anatomical regions evolved in response to directional selection, stabilizing selection, neutral evolutionary forces, or a combination of the three among taxa. It likewise remains unknown if function was a motivating selection factor in shaping both trait covariances and evolution. The ability of traits to respond to directional selection is dependent on both the covariance structure among traits, and the strength of those connections (see Rolian, 2014). Both of these factors were shown to be related to functional differences among the primate hands and feet (Rolian, 2009) and limbs (Young et al., 2010). However, studies of the axial skeleton indicate similar covariance structures among the face and neurocranium of Catarrhines (de Oliveira et al., 2009), and the pelvis (Lewton, 2012) and scapula (Young, 2004) across primates. For the skull and scapula, the strength of the covariance among traits appears to be more influential on the morphological variation among traits. The comparisons across primate taxa performed in this study support these findings. This result was somewhat unexpected, as there is a large degree of morphological variation in the primate shoulder girdle that is reflective of differences in the functional demands of the upper limb (Inman et al., 1944; Ashton and Oxnard, 1961, 1963; Oxnard, 1963; Roberts, 1974; Stern et al., 1977; Tuttle and Basmajian, 1977, 1978; Jungers and Stern, 1984; Larson and Stern, 1986; Larson et al., 1991; Larson, 1993, 1998; Whitehead and Larson, 1994; Preuschoft et al., 2010). Traits with direct connection to the shoulder girdle are all participating in the same basic function: movement of the shoulder and upper limb. Other studies have attributed the maintenance of the covariance structure among traits to not be a remnant of phylogeny, but rather internal stabilizing selection on the determinants of the covariance structure (Marroig and Cheverud, 2001).

Therefore, we conclude that while our study provides abundant novel evidence about the patterns of constraint and potential evolution amongst the four anatomical regions we consider, our results also show a wealth of future avenues of inquiry. These could clarify how traits within anatomical regions contribute to evolutionary constraint across regions, and thus the whole organism, as well as the genetic, developmental, and functional factors that underlie these evolutionary relationships. Furthermore, our analysis indicates that the ways in which we standardize data is a non-trivial concern in pursuing these types of questions. As we show, mean standardization magnifies the influence of differences in organism size, which can overwhelm more subtle within organism effects when comparing trait evolvabilities (compare, for example, evolutionary flexibility in Table 3 and SOM Table S8). While this is not a novel observation (see for example Hansen and Houle, 2008; Rolian 2009; Houle et al., 2011) our analyses highlight its importance.

## 5. Conclusions

This study demonstrates evolutionary non-independence of the primate shoulder girdle, as its evolutionary potential is affected by its covariance with other anatomical regions included in this study, namely the basicranium, pelvis, and humerus. Our findings add further evidence for broadly shared patterns of potential evolutionary constraint across multiple primate taxa. While evolutionary covariance is observed among these anatomical regions for all taxa, the patterns of the strength of these covariances differ. The patterns of morphological integration among anatomical regions are remarkably similar among primate genera, whereas the patterns of evolutionary flexibility among genera vary. The evolutionary potential does not follow a pattern related to functional variation in the upper limb, rather it more closely reflects a dichotomy between monkeys and apes, with apes generally exhibiting less potential evolutionary constraint among anatomical regions. These results suggest the evolutionary and thus underlying genetic relationships among anatomical regions differ among primate genera, and those differences may have contributed to the morphological diversity observed among extant primate taxa. While this study elucidates a previously uninvestigated factor affecting primate morphological variation, these results can only assess the evolutionary potential among traits in response to directional selection. Future analyses using different approaches will be necessary to discern by which evolutionary processes shaped trait variation, and the effect the reported patterns of morphological integration and evolutionary flexibility among anatomical regions and between primate genera, respectively, had on their morphological evolution.

## Supporting information

Agosto_Auerbach_Primate_Flexibility_SOM_2022

## Acknowledgments

We thank C. Roseman, B. O’Meara, and G. Cabana for helpful comments on an early draft of this paper. E.R.A. thanks the many North American institutions for access to collections used for this study. This research was supported by grants to B.M.A and E.R.A. from the National Science Foundation (Doctoral Dissertation Improvement Grant BCS–1825995), and a Thomas Fellowship through the University of Tennessee and the College of Arts and Sciences at the University of Tennessee to E.R.A.

