## Supplementary material for "Morphological integration and evolutionary potential of the primate shoulder: Variation among taxa and implications for genetic covariances with the basicranium, pelvis, and arm": Agosto_Auerbach_Primate_Flexibility_SOM_2022

### SOM S1

#### Combining species for *Pongo* and *Papio*

Lack of preservation of the crania and os coxa prevented the measurement of several traits in these regions for most taxa, with *Pongo* and *Papio* being the most heavily affected. Missing values could not be estimated because of the non-random nature of missing measurements (Little and Rubin, 2019). Rather, the samples for both *Pongo* and *Papio* are comprised of pooled data from multiple species to achieve the minimum sample sizes recommended by Grabowski and Porto (2017) for estimating evolutionary flexibility ($n \geq 11$ for 10 traits, via the 'howmany' R function; see SOM Table S1). Pooling species for these samples was deemed reasonable based on nonsignificant ($p > 0.05$) differences in the measures used, as indicated by a multivariate analysis of variance (see SOM Table S2). Moreover, differences among trait covariances are expected to be higher between genera than among closely related species of the same genus (de Oliveira et al., 2009).

#### Noise Correction

In the calculation of both conditioned covariance and trait autonomy, the *P* matrix is inverted to allow for matrix multiplication. When a matrix is inverted, the eigenvectors are likewise inverted, and so the smallest eigenvectors will have the greatest values. As our matrices consist of 20 traits, there are 20 eigenvectors, and not all of these will encompass a high percentage of the variance. As noted by Twede and Hayden (2004), the smallest eigenvectors in a matrix will often be more subject to including random residual variance (i.e., 'noise'), especially variance that is introduced by measurement error. For this reason, the effect of the smallest eigenvectors on the inverted *P* matrix must be minimized.

One solution, borne from methods to remove pixilation and static from images, is to replace eigenvalues of the smallest eigenvectors with the median value (Twede and Hayden, 2004; Marroig et al., 2012). We controlled for noise in our variance-covariance (VCV) matrix estimates by replacing values smaller than the median of the variance of the second derivative derived from the absolute value of eigenvalues with the median, and recalculating the new VCV matrix using these new eigenvalues (Twede and Hayden, 2004). All analyses were performed using the noise-corrected VCV matrices.

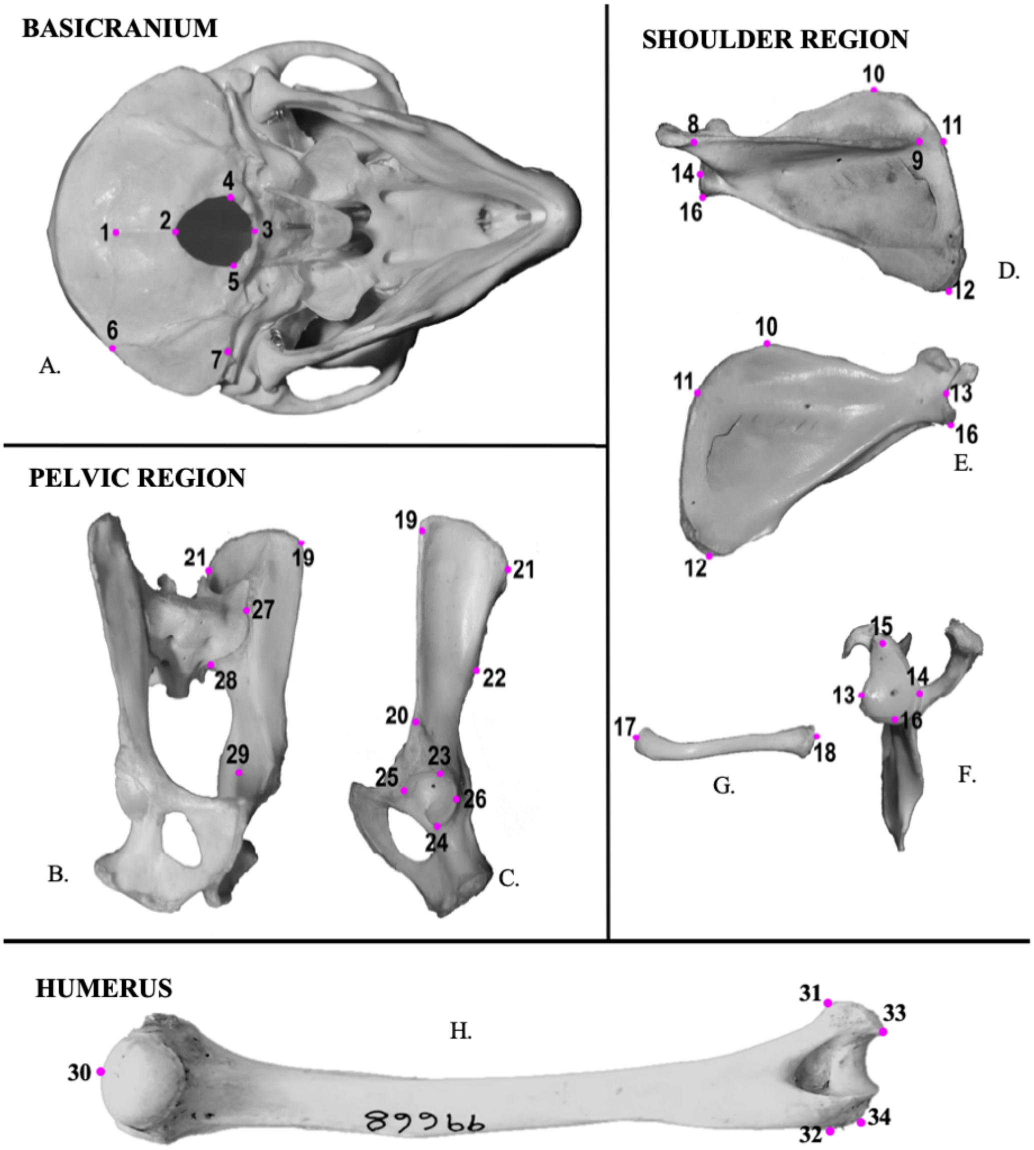

**SOM Figure S1.**

Landmarks taken from each anatomical region: A) caudal aspect of the basicranium; B) medial aspect of os coxa; C) lateral aspect of os coxa; D) dorsal aspect of scapula; E) ventral aspect of scapula; F) lateral aspect of scapula; G) cranial aspect of clavicle; H) ventral aspect of humerus. Refer to SOM Table S4 for a list of landmarks and descriptions. Landmark locations are demonstrated on a *Macaca mulatta* specimen and are homologous for all tax

***A. Alouatta***

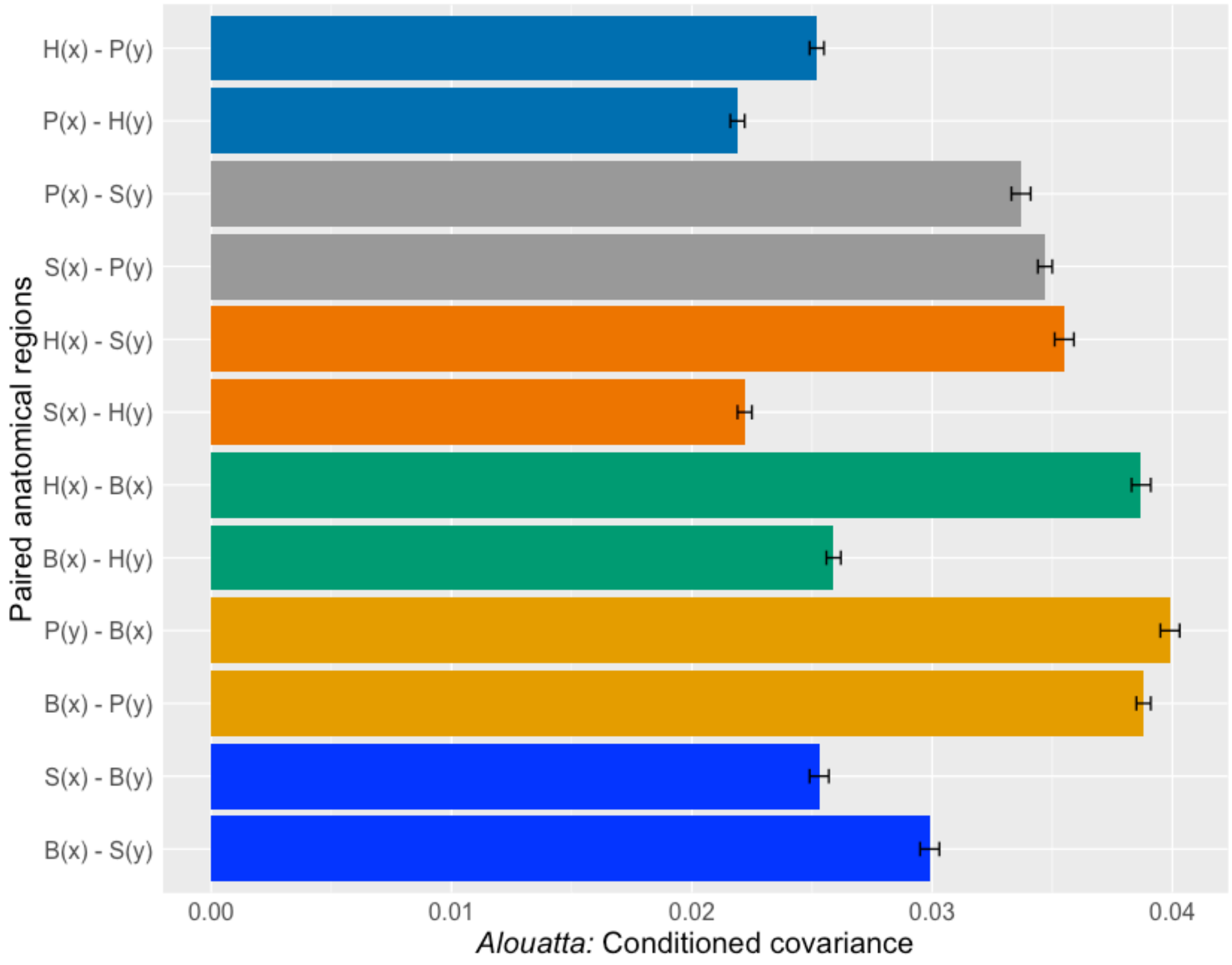

**SOM Figure S2.**

Conditioned covariance and 95% confidence intervals using size corrected variance covariance matrices for each genus included in this study: A) *Alouatta*, B) *Colobus*, C) *Macaca*, D) *Papio*, E) *Homo*, F) *Hylobates*, G) *Pan* and H) *Pongo*. Estimates of conditioned covariance are compared between pairings of the following anatomical regions: basicranium (B), shoulder girdle (S), pelvic girdle (P) and humerus (H). Conditioned covariance is calculated twice for each pairing, with anatomical region standing in for the traits under stabilizing selection (x) and the traits responding to directional selection (y).

***B. Colobus***

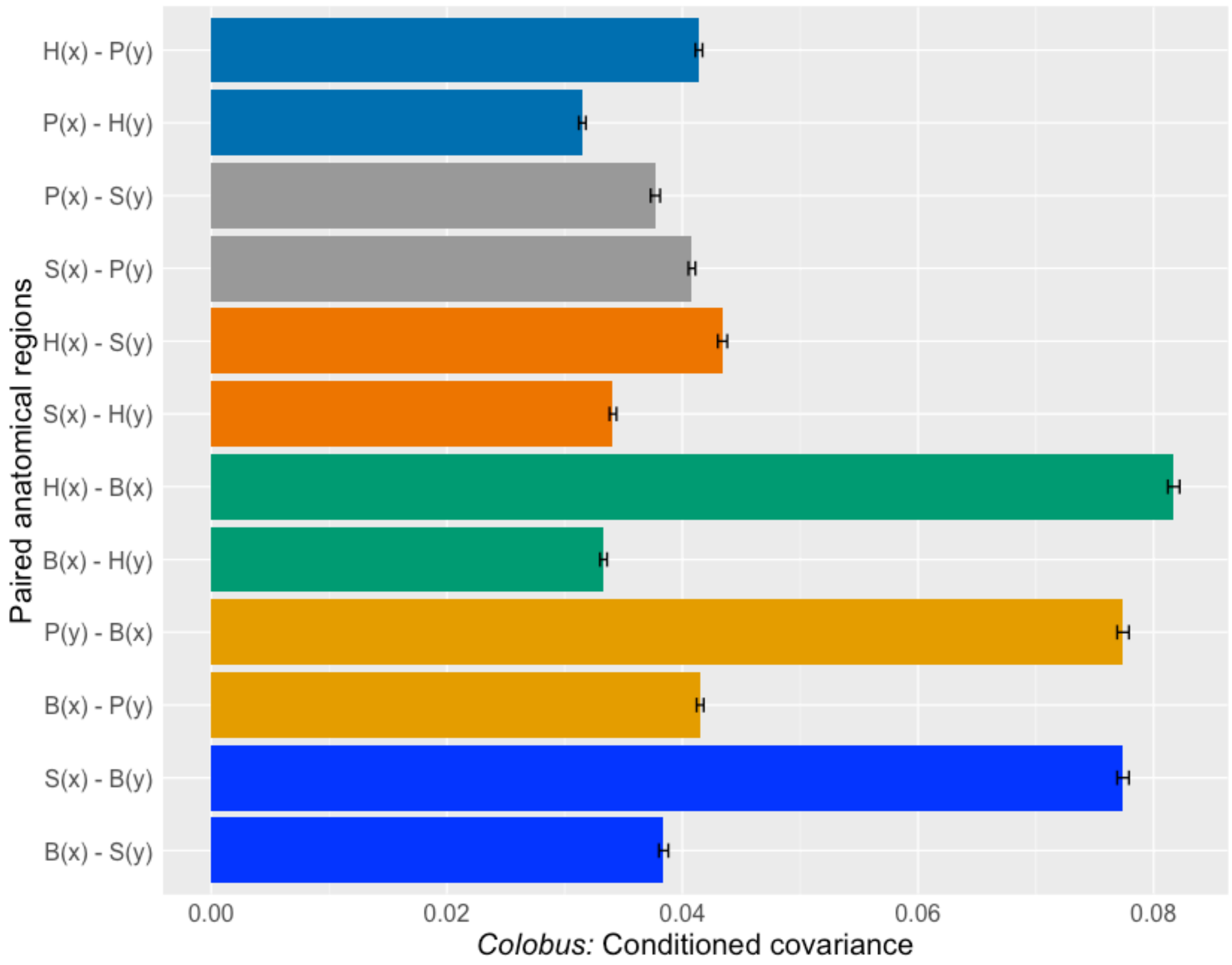
H(x) - P(y)
P(x) - H(y)
P(x) - S(y)
S(x) - P(y)
H(x) - S(y)
S(x) - H(y)
H(x) - B(x)
B(x) - H(y)
P(y) - B(x)
B(x) - P(y)
S(x) - B(y)
B(x) - S(y)
Paired anatomical regions
0.00
0.02
0.04
0.06
0.08
*Colobus:* Conditioned covariance

***C. Macaca***

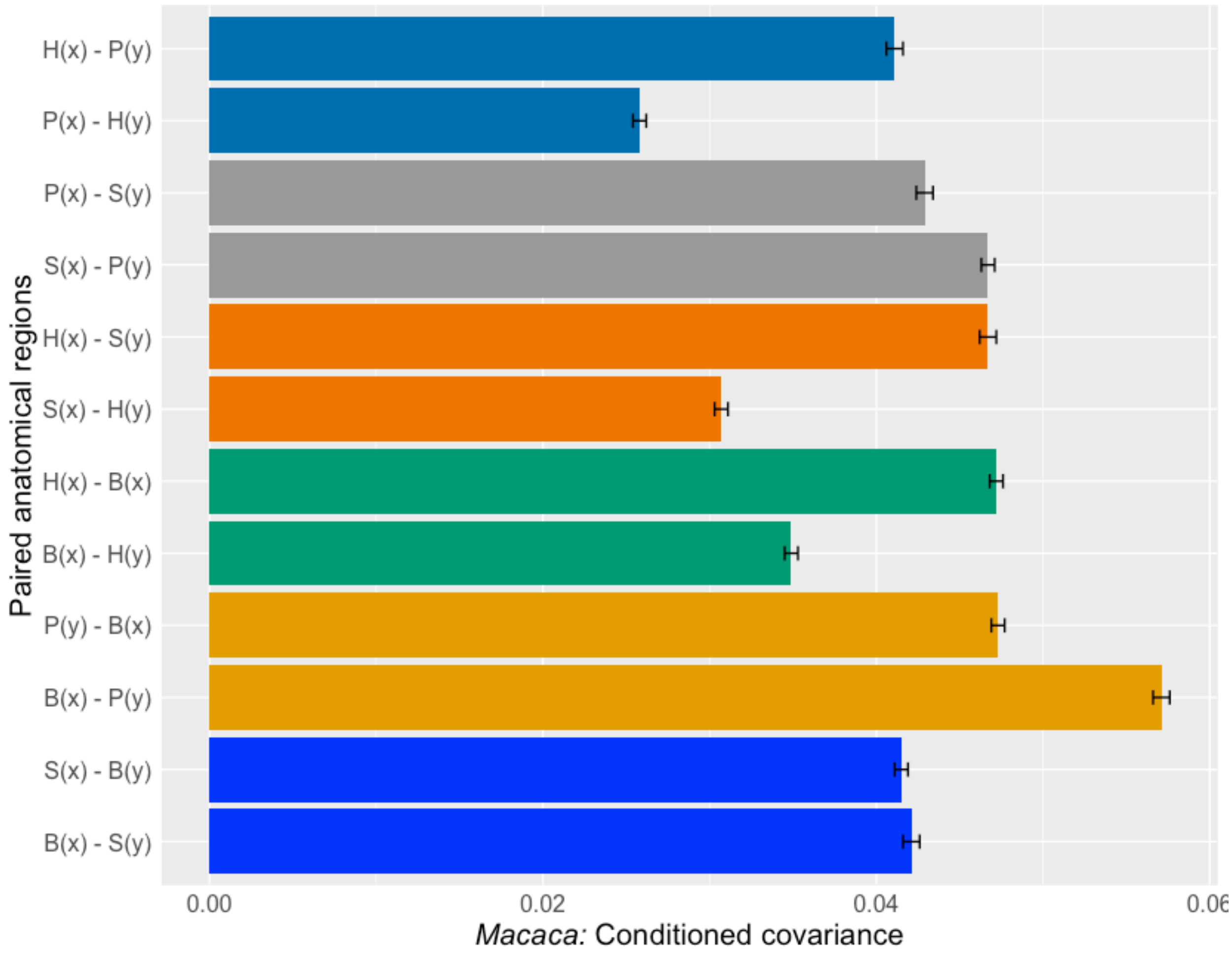
H(x) - P(y)
P(x) - H(y)
P(x) - S(y)
S(x) - P(y)
H(x) - S(y)
S(x) - H(y)
H(x) - B(x)
B(x) - H(y)
P(y) - B(x)
B(x) - P(y)
S(x) - B(y)
B(x) - S(y)
Paired anatomical regions
0.00
0.02
0.04
0.06
*Macaca:* Conditioned covariance

***D. Papio***

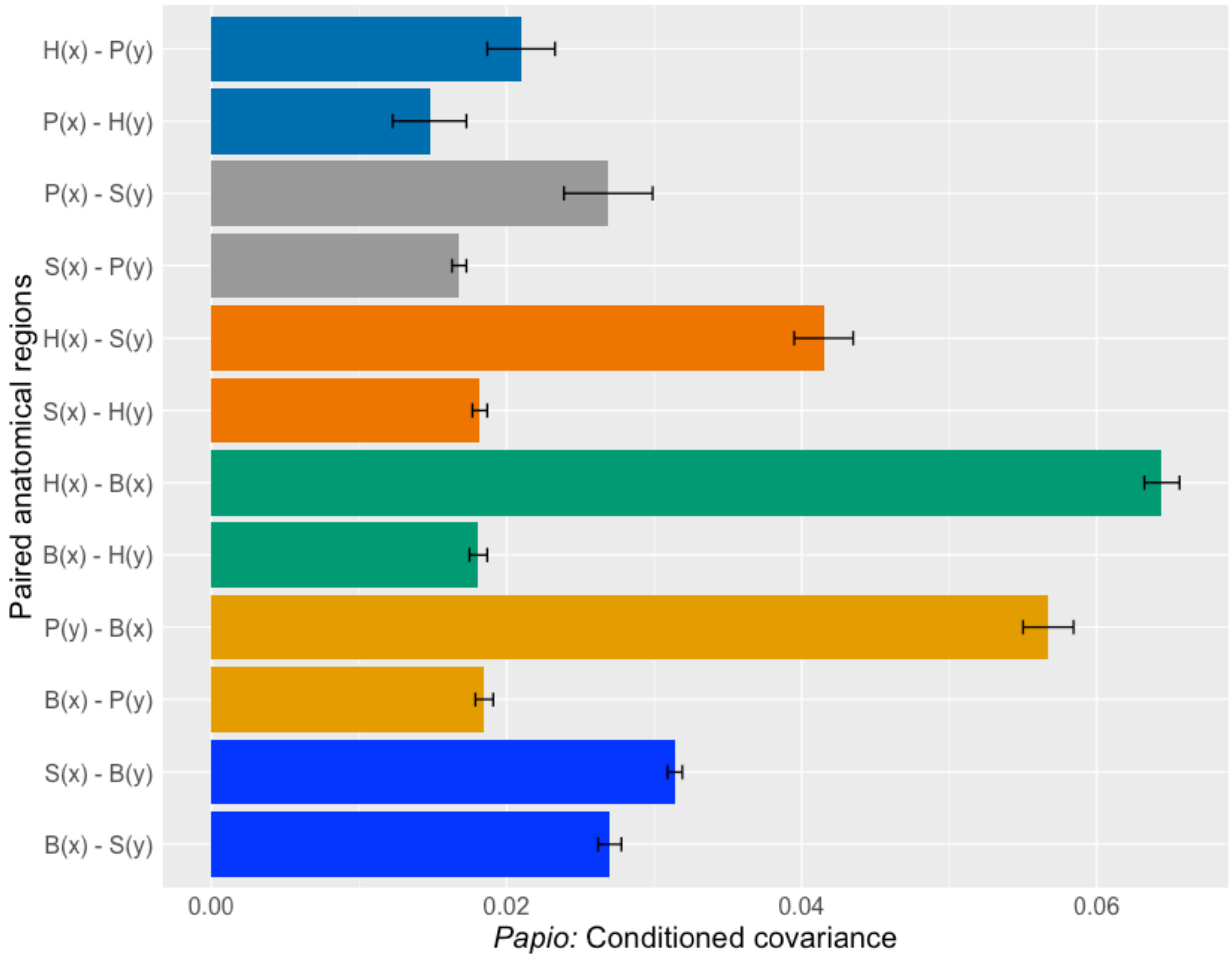
H(x) - P(y)
P(x) - H(y)
P(x) - S(y)
S(x) - P(y)
H(x) - S(y)
S(x) - H(y)
H(x) - B(x)
B(x) - H(y)
P(y) - B(x)
B(x) - P(y)
S(x) - B(y)
B(x) - S(y)
Paired anatomical regions
0.00
0.02
0.04
0.06
Papio: Conditioned covariance

***E. Homo***

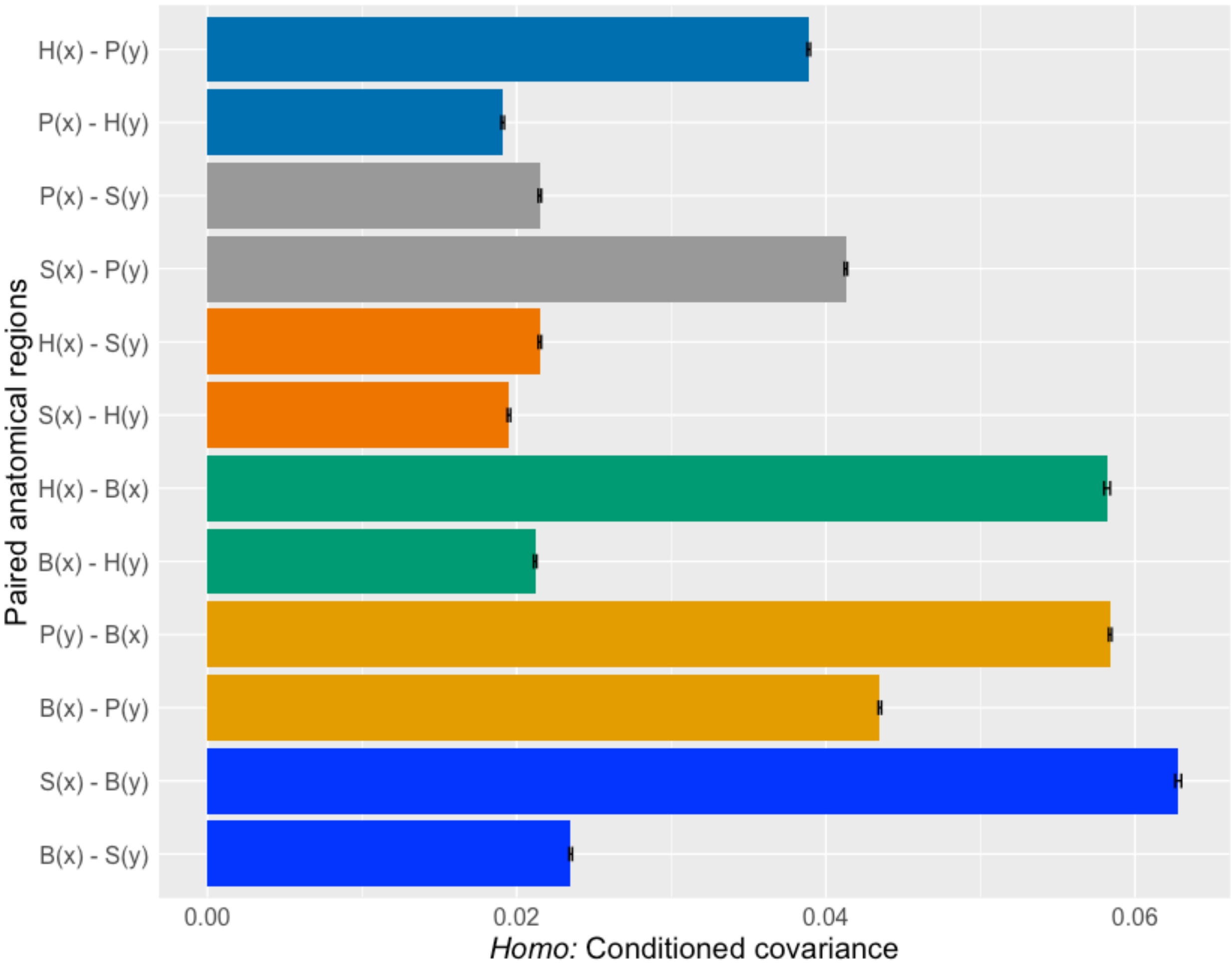

H(x) - P(y)
P(x) - H(y)
P(x) - S(y)
S(x) - P(y)
H(x) - S(y)
S(x) - H(y)
H(x) - B(x)
B(x) - H(y)
P(y) - B(x)
B(x) - P(y)
S(x) - B(y)
B(x) - S(y)
Paired anatomical regions
0.00
0.02
0.04
0.06
*Homo:* Conditioned covariance

***F. Hylobates***

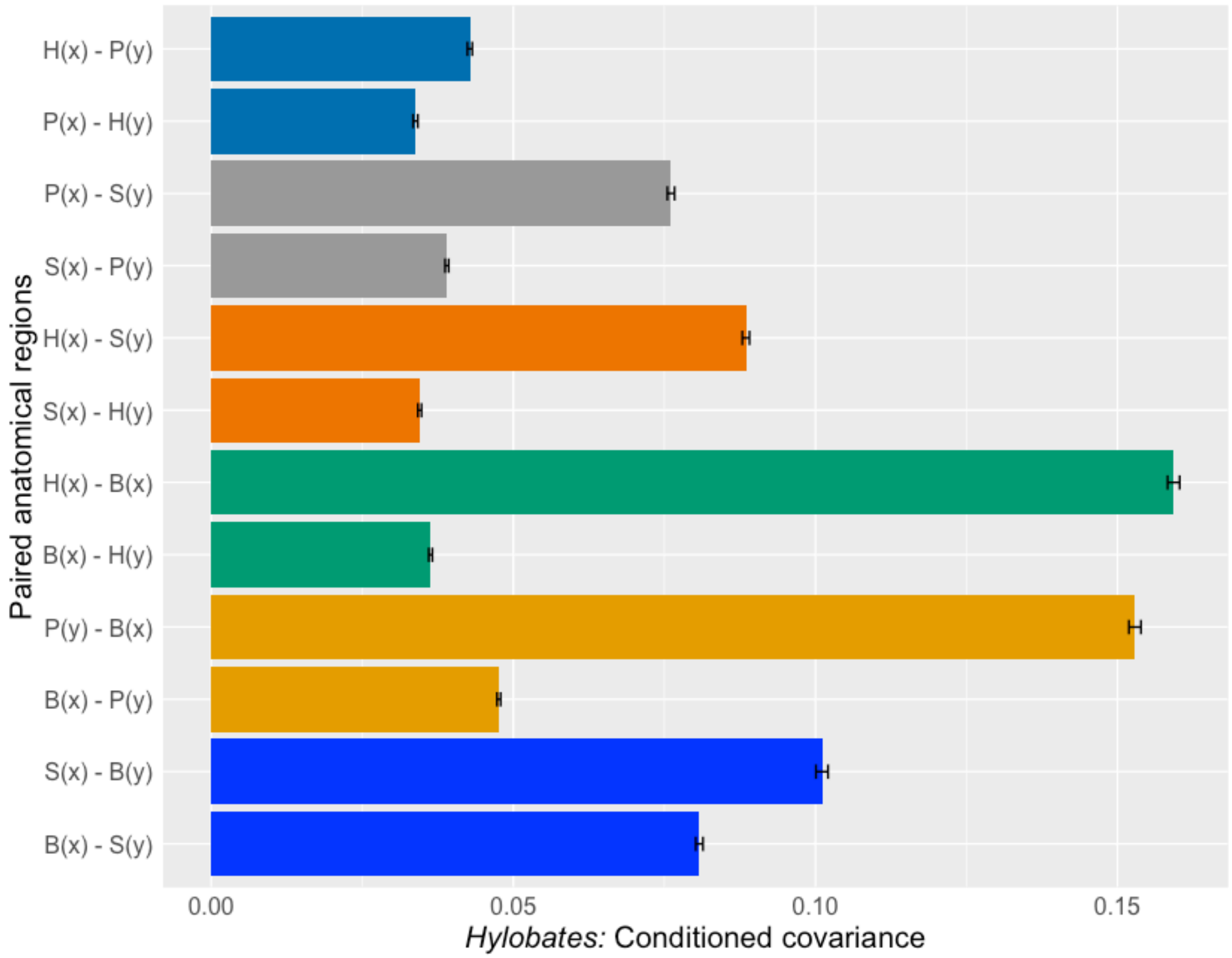

H(x) - P(y)
P(x) - H(y)
P(x) - S(y)
S(x) - P(y)
H(x) - S(y)
S(x) - H(y)
H(x) - B(x)
B(x) - H(y)
P(y) - B(x)
B(x) - P(y)
S(x) - B(y)
B(x) - S(y)
Paired anatomical regions
0.00
0.05
0.10
0.15
*Hylobates:* Conditioned covariance

*G. Pan*

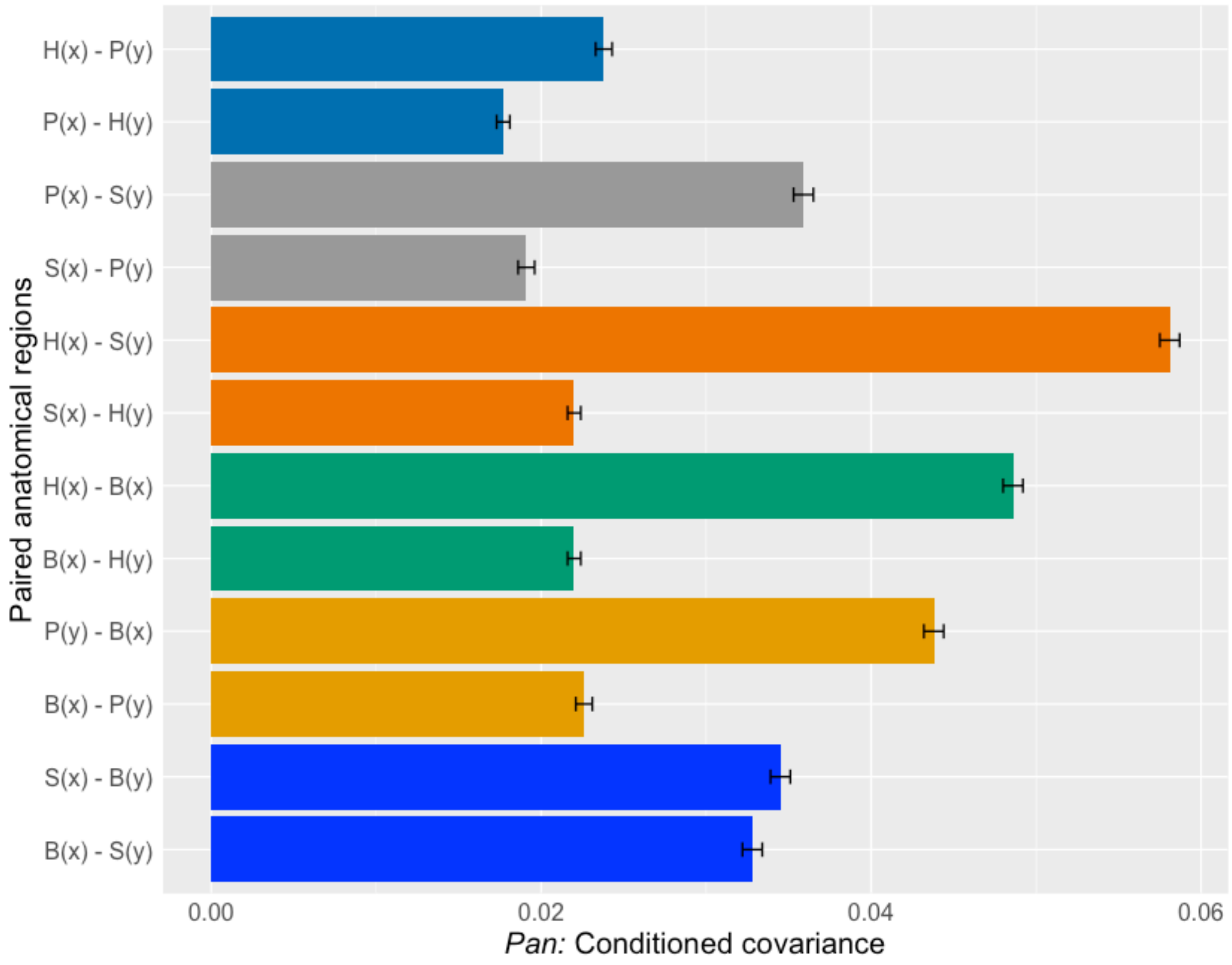

H(x) - P(y)
P(x) - H(y)
P(x) - S(y)
S(x) - P(y)
H(x) - S(y)
S(x) - H(y)
H(x) - B(x)
B(x) - H(y)
P(y) - B(x)
B(x) - P(y)
S(x) - B(y)
B(x) - S(y)
Paired anatomical regions
0.00
0.02
0.04
0.06
*Pan:* Conditioned covariance

***H. Pongo***

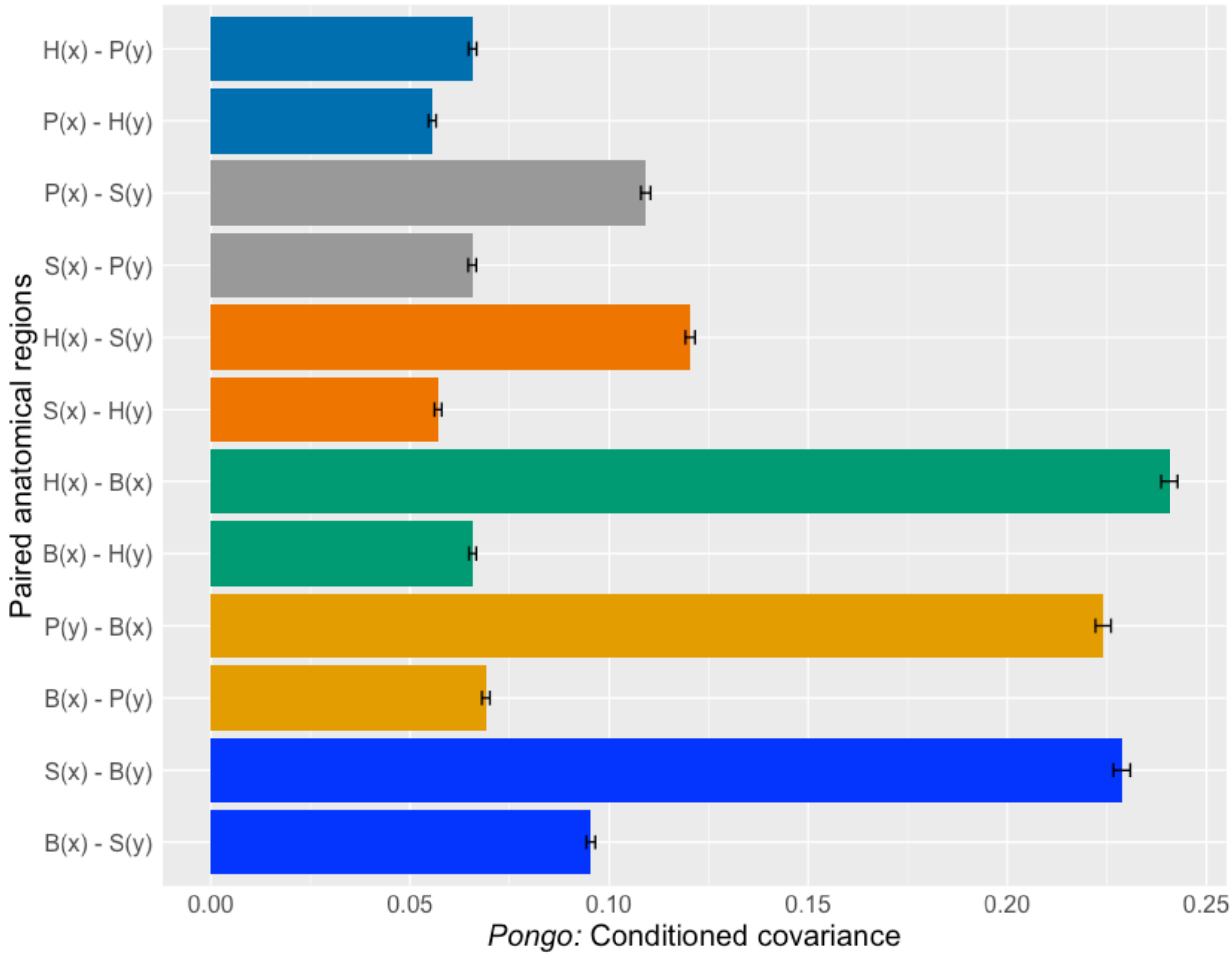
H(x) - P(y)
P(x) - H(y)
P(x) - S(y)
S(x) - P(y)
H(x) - S(y)
S(x) - H(y)
H(x) - B(x)
B(x) - H(y)
P(y) - B(x)
B(x) - P(y)
S(x) - B(y)
B(x) - S(y)
Paired anatomical regions
0.00
0.05
0.10
0.15
0.20
0.25
Pongo: Conditioned covariance

**SOM Table S1**

Results of the 'howmany' R function (Grabowski and Porto, 2017) to estimate minimum recommended sample size for evolvability statistics using the recommended MI (R2) range of 0.04 (left) to 0.5 (right) for both 10 (top) and 20 (bottom) traits.

| **Ac = 0.05, MI = 0.04, n trait = 10** | |
|---|---|
| **Statistic** | **Individuals** |
| Respondability | 34.09043 |
| Evolvability | 11 |
| Conditional evolvability | 64.53546 |
| Integration | 29.76976 |
| R2 | 117.05921 |
| Flexibility | 11 |

| **Ac = 0.05, MI = 0.5, *n* trait = 10** | |
|---|---|
| **Statistic** | **Individuals** |
| Respondability | 110.46316 |
| Evolvability | 35.96475 |
| Conditional evolvability | 64.53546 |
| Integration | 11.05206 |
| R2 | 17.80881 |
| Flexibility | 11 |

| **Ac = 0.05, MI = 0.04, *n* trait = 20** | |
|---|---|
| **Statistic** | **Individuals** |
| Respondability | 35.97704 |
| Evolvability | 21 |
| Conditional evolvability | 98.27549 |
| Integration | 23.59836 |
| R2 | 117.05921 |
| Flexibility | 21 |

| **Ac = 0.05, MI = 0.5, *n* trait = 20** | |
|---|---|
| **Statistic** | **Individuals** |
| Respondability | 81.8756 |
| Evolvability | 36.8137 |
| Conditional evolvability | 98.27549 |
| Integration | 21 |
| R2 | 21 |
| Flexibility | 21 |

Abbreviations: Ac = maximum inaccuracy, MI = overall level of integration, *n* trait = number of traits within the analysis, R2 = average of squared correlations among traits.

**SOM Table S2**

Results of multivariate analysis of variance between the sexes and among species of baboons (*Papio*) and orangutans (*Pongo*).

| | | | | | | |
|---|---|---|---|---|---|---|
| | | | ***Papio*** | | | |
| **Analysis** | | **Df** | **Sum squares** | **Mean squares** | **F-value** | ***p*-value[a]** |
| | Sex | 1 | 401222 | 401222 | 34.653 | **p < 0.001** |
| Sex * species | Species | 2 | 79012 | 39506 | 3.412 | 0.060 |
| | Sex:species | 1 | 136 | 136 | 0.012 | 0.915 |
| | Residuals | 15 | 173674 | 11578 | | |
| | | | ***Pongo*** | | | |
| **Analysis** | | **Df** | **Sum squares** | **Mean squares** | **F-value** | ***p*-value** |
| | Sex | 1 | 602000 | 602000 | 41.429 | **p < 0.001** |
| Sex * species | Species | 2 | 9636 | 4818 | 0.332 | 0.721 |
| | Residuals | 21 | 305150 | 14531 | | |

[a]Bolded values are statistically significant at $p < 0.05$.

### SOM Table S3

Trait abbreviations, definitions and their functional and/or developmental correlates for each anatomical region.[a]

| Trait abbreviation | Trait definition |
|---|---|
| | **Basicranium** |
| **NR** | Length of the bony attachment of the nuchal ligament on the basicranium (Richmond et al., 2001). The nuchal ligament serves as an origin for the trapezius muscle (Diogo and Wood, 2012), and has partial contributions from post-otic neural crest (PONC) during development (Matsuoka et al., 2005). |
| **SCM** | Length of cranial insertion of the sternocleidomastoid mastoid muscle (Richmond et al., 2001). This muscle and its attachments have PONC contributions during development (Matsuoka et al., 2005). |
| **SNL** | Length of the caudal border of the attachment for cranial-most neck muscles and trapezius muscle (Richmond et al., 2001). The attachment of the trapezius has PONC contribution during development (Matsuoka et al., 2005). |
| **FMW** | Width of the foramen magnum measured at the greatest curvature of the left and right lateral margins (Moore-Jansen et al., 1994). This provides a measure of the intercondylar breadth for the occipital condyles, which are derived from the somites during ontogeny (Pilbeam, 2004). |
| **FML** | Ventral to dorsal length of the foramen magnum (Moore-Jansen et al., 1994). Derived from paraxial mesoderm and somites during development (Pilbeam, 2004). |
| | **Shoulder Girdle** |
| **SSL** | Length of the scapular spine—insertion of the trapezius muscle along the scapular spine (Richmond et al., 2001; Young, 2006). This point of attachment is derived from PONC during development (Matsuoka et al., 2005). |
| **SBL** | Length of scapular blade along lateral border (Martin, 1928). Combined origin length for teres minor, teres major, latissimus dorsi (in some primates), and long head of the triceps brachii muscles (Diogo and Wood, 2012; Zihlman and Underwood, 2019). |
| **SBW** | Maximum medial to lateral breadth of scapular blade (Martin, 1928; Moore-Jansen et al., 1994). Roughly corresponds to medial-lateral length of the caudal portion of supraspinatus and cranial portion of infraspinatus muscles. |
| **SBH** | Cranial to caudal height of scapular blade (Martin, 1928; Moore-Jansen et al., 1994). Combined length of the supraspinous and infraspinous fossae, which provide the combined breadth of the supraspinatus and infraspinatus muscles (Diogo and Wood, 2012; Zihlman and Underwood, 2019). |
| **VB** | Length of the vertebral border of the scapula. Combined values of VBA (distance between landmarks 10-11) and VBB (distance between landmarks 11-12). The vertebral border of the scapula is the location of the insertion of rhomboid major and rhomboid minor muscles (Diogo and Wood, 2012; Zihlman and Underwood, 2019), and is derived from the somites during development (Valasek et al., 2010). |

| Trait Abbreviation | Trait definition |
|---|---|
| **GW** | Maximum ventral to dorsal width of glenoid fossa (Young, 2006). Maximum width of the shoulder joint surface, derived from the somatopleure during development (Huang et al., 2000, 2006; Sears et al., 2015). |
| **GL** | Maximum cranial to caudal length of the glenoid fossa (Young, 2006). Maximum height of the shoulder joint surface, derived from the somatopleure during development (Huang et al., 2000, 2006; Sears et al., 2015). |
| **CLML** | Maximum length of the clavicle (Martin, 1928; Moore-Jansen et al., 1994). Part of the bony shoulder girdle, and medial portion is the origin for the sternocleidomastoid muscle and is derived from PONC during development (Matsuoka et al., 2005). |
| | **Pelvic Girdle** |
| **AMIB** | Anterior margin of iliac blade (Grabowski and Roseman, 2015). Distance is functionally related to positioning of the lesser gluteal muscles which can aide in pelvic stabilization during the single support phase in bipedal locomotion (Sigmon, 1975). Analogous blade measurement to the lateral border of the scapula. |
| **PMIB** | Posterior margin of iliac blade (Grabowski and Roseman, 2015). Shorter distances place the body's center of mass above the hips during more bipedal locomotion (see McCollum et al., 2010). Analogous to the medial blade border of scapula. |
| **ASL** | Cranial to caudal length of the auricular surface (Grabowski and Roseman, 2015). Functionally related to weight bearing (Berge and Kazmierczak, 1986). |
| **RAH** | Retro-auricular height, distance from cranial margin of the auricle to posterior superior iliac spine (Grabowski and Roseman, 2015). Expansion of this area is related to more upright trunk postures (see Lovejoy, 2005; Stern and Susman, 1983; Bramble and Lieberman, 2004). Roughly analogous to scapular blade breadth. |
| **LIB** | Lateral iliac breadth, distance from cranial margin of the auricle to the anterior superior iliac spine (Grabowski and Roseman, 2015). Roughly analogous to scapular blade breadth. |
| **LIH** | Lower iliac height, distance from caudal margin of auricle to the posterior aspect of the central point of the acetabulum (Grabowski and Roseman, 2015). Longer distances related to lumbar mobility (see Lovejoy et al., 2009). Analogous to scapular blade length. |
| **AL** | Cranial to caudal length of the acetabulum (Lewton, 2012). Analogous measure to GL as part of the joint surface. |
| **AW** | Medial to lateral width of the acetabulum. Is an analogous measure to GW as part of the joint surface, and can be used as a proxy for body size (Ruff, 2003). |
| | **Humerus** |
| **HML** | Maximum length of the humerus (Moore-Jansen et al., 1994). |
| **MCH** | Length of the bony attachment of several flexor muscles of the forearm (Zihlman and Underwood, 2019). |
| **LCH** | Length of the bony attachment of several extensor muscles of the forearm (Zihlman and Underwood, 2019). |

| Trait Abbreviation | Trait definition |
|---|---|
| **HEW** | Medial to lateral width of distal humerus at the epicondyles (Zihlman and Underwood, 2019). |
| **HDW** | Medial to lateral width of the distal articular region of the humerus (Zhilmand and Underwood, 2019). Articular surface of elbow joint. |

[a]Definitions are modified from Agosto and Auerbach (2021).

**SOM Table S4**

Definitions of landmarks collected. Landmark numbers correspond to labeled landmark locations in SOM Figure S1.

| No. | Landmark | Definition |
|---|---|---|
| | | Basicranium |
| 1. | Inion | Location of the intersection of the right and left superior nuchal lines within the external occipital protuberance (White et al., 2012). |
| 2. | Opisthion | Midline point of the dorsal margin of the foramen magnum (White et al., 2012). |
| 3. | Basion | Midline point of the ventral margin of the foramen magnum (White et al., 2012). |
| 4. | Right margin of foramen magnum | Right margin of the foramen magnum at its greatest curvature (Moore-Jansen et al., 1994). |
| 5. | Left margin of foramen magnum | Left margin of the foramen magnum at its greatest curvature (Moore-Jansen et al., 1994). |
| 6. | Asterion | Point where the lambdoidal, parietomastoid and occipitomastoid sutures meet (White et al., 2012). |
| 7. | Left mastoid | Inferior-most point of the mastoid process (White et al., 2012). |
| | | Scapula |
| 8. | Scapular spine lateral border | Point of intersection between the scapular spine and acromion process (Sears et al., 2013). |
| 9. | Scapular spine medial border | Medial margin of trapezius attachment along scapular spine (Young, 2006). |
| 10. | Superior angle | Intersection of superior and medial borders of scapula (White et al., 2012). |
| 11. | Scapular spine vertebral border | Point on vertebral margin where the long axis of the scapular spine and vertebral border meet (Young, 2006). |
| 12. | Inferior angle | Intersection of the lateral and medial borders of scapula (White et al., 2012). |
| 13. | Ventral margin of glenoid fossa | Ventral margin of glenoid fossa at its greatest width (Young, 2006). |
| 14. | Dorsal margin of glenoid fossa | Dorsal margin of glenoid fossa at its greatest width (Young, 2006). |
| 15. | Cranial margin of glenoid fossa | Superior-most margin of glenoid fossa (Young, 2006). |
| 16. | Caudal margin of glenoid fossa | Inferior-most margin of glenoid fossa (Young, 2006). |
| | | Clavicle |
| 17. | Acromial end | Lateral-most point of clavicle on cranial aspect. |
| 18. | Sternal end | Medial-most point of clavicle on cranial aspect. |

| No. | Landmark | Definition |
|---|---|---|
| | | Os coxa |
| 19. | Anterior superior iliac spine | Ventral-most point on lateral aspect of iliac crest (Lewton, 2012). |
| 20. | Anterior inferior iliac spine | Ventral-most point on anterior inferior iliac spine. If only a bony roughening, the point is taken at the center of this rugosity (Lewton, 2012). |
| 21. | Posterior superior iliac spine | Medial-most point of the dorsal-cranial border of iliac crest (Lewton, 2012). |
| 22. | Posterior inferior iliac spine | Sharp projection posterior and inferior to the auricular surface (White et al., 2012). |
| 23. | Cranial margin of acetabulum | Cranial-most point of the margin of the acetabulum (Lewton, 2012). |
| 24. | Caudal margin of acetabulum | Caudal-most point of the margin of the acetabulum, directly across from landmark 23 (Lewton, 2012). |
| 25. | Medial margin of acetabulum | Medial-most point along margin of the acetabulum. |
| 26. | Lateral margin of acetabulum | Lateral-most point along margin of the acetabulum. |
| 27. | Cranial margin of auricular surface | Central point of the cranial-most margin of the auricular surface. |
| 28. | Caudal margin of auricular surface | Central point of the caudal-most margin of the auricular surface. |
| 29. | Center of acetabulum | Center of the acetabulum, defined as the intersection of the line between landmarks 23 and 24 and landmarks 27 and 28 (Lewton, 2012). |
| | | Humerus |
| 30. | Cranial end of humerus | Cranial-most point of the proximal end of humerus. |
| 31. | Medial epicondyle | Medial-most margin of medial epicondyle of humerus. |
| 32. | Lateral epicondyle | Lateral-most margin of lateral epicondyle of humerus. |
| 33. | Medial margin of distal articular surface | Medial-most margin of distal articular surface of the humerus. |
| 34. | Lateral margin of distal articular surface | Lateral-most margin of distal articular surface of the humerus. |

**SOM Table S5**

Estimates of evolvability, standard error (indicated by ±), and 95% confidence intervals (in parentheses) between pairs of anatomical regions for each genus using size corrected variance covariance matrices.

| | Taxon | B-P | B-S | S-P | S-H | B-H | P-H |
|---|---|---|---|---|---|---|---|
| **Platyrrhines** | ***Alouatta*** | 0.0108 ± < 0.0001 (0.0107, 0.0109) | 0.0112 ± < 0.0001 (0.0110, 0.0113) | 0.0118 ± < 0.0001 (0.0117, 0.0119) | 0.0102 ± < 0.0001 (0.01, 0.0103) | 0.0089 ± < 0.0001 (0.0088, 0.009) | 0.0097 ± < 0.0001 (0.0096, 0.0098) |
| **Cercopithecoids** | ***Colobus*** | 0.0134 ± < 0.0001 (0.0133, 0.0136) | 0.0132 ± < 0.0001 (0.0131, 0.0134) | 0.0091 ± < 0.0001 (0.009, 0.0093) | 0.0079 ± < 0.0001 (0.0078, 0.008) | 0.0122 ± < 0.0001 (0.0121, 0.0124) | 0.0082 ± < 0.0001 (0.0081, 0.0083) |
| | ***Macaca*** | 0.0221 ± < 0.0001 (0.0219, 0.0224) | 0.0272 ± < 0.0001 (0.027, 0.0275) | 0.0156 ± < 0.0001 (0.0154, 0.0158) | 0.0211 ± < 0.0001 (0.0209, 0.0213) | 0.0093 ± < 0.0001 (0.0091, 0.0094) | 0.0147 ± < 0.0001 (0.0145, 0.0148) |
| | ***Papio*** | 0.0264 ± 0.0004 (0.0256, 0.0273) | 0.0382 ± 0.0005 (0.0372, 0.0392) | 0.0353 ± 0.0005 (0.0342, 0.0363) | 0.0318 ± 0.0006 (0.0307, 0.0329) | 0.0228 ± 0.0005 (0.0219, 0.0237) | 0.0197 ± 0.0005 (0.0188, 0.0206) |
| **Hominoids** | ***Homo*** | 0.0057 ± < 0.0001 (0.0056, 0.0057) | 0.0064 ± < 0.0001 (0.0063, 0.0065) | 0.0059 ± < 0.0001 (0.0058, 0.006) | 0.0046 ± < 0.0001 (0.0046, 0.0047) | 0.0044 ± < 0.0001 (0.0043, 0.0045) | 0.0037 ± < 0.0001 (0.0037, 0.0038) |
| | ***Hylobates*** | 0.0221 ± 0.0001 (0.0219, 0.0224) | 0.0272 ± 0.0001 (0.027, 0.0275) | 0.0156 ± < 0.0001 (0.0154, 0.0158) | 0.0147 ± < 0.0001 (0.0145, 0.0148) | 0.0211 ± 0.0001 (0.0209, 0.0213) | 0.0093 ± < 0.0001 (0.0091, 0.0094) |
| | ***Pan*** | 0.0092 ± < 0.0001 (0.009, 0.0093) | 0.0131 ± < 0.0001 (0.013, 0.0133) | 0.0108 ± < 0.0001 (0.0106, 0.011) | 0.0101 ± < 0.0001 (0.0099, 0.0103) | 0.0083 ± < 0.0001 (0.0082, 0.0085) | 0.0062 ± < 0.0001 (0.006, 0.0063) |
| | ***Pongo*** | 0.0368 ± 0.0003 (0.0363, 0.0374) | 0.0436 ± 0.0003 (0.0431, 0.0441) | 0.0238 ± 0.0003 (0.0234, 0.0241) | 0.0225 ± 0.0003 (0.0221, 0.0228) | 0.0357 ± 0.0003 (0.0352, 0.0363) | 0.0159 ± 0.0002 (0.0155, 0.0162) |

Abbreviations: B-P = basicranium and pelvic girdle, B-S = basicranium and shoulder girdle, S-P = shoulder girdle and pelvic girdle, S-H = shoulder girdle and humerus, B-H = basicranium and humerus, P-H = pelvic girdle and humerus.

**SOM Table S6**

Estimates of evolvability, standard error (indicated by ±), and 95% confidence intervals (in parentheses) between pairs of anatomical regions for each genus using mean standardized variance covariance matrices.

| | Taxon | B-P | B-S | S-P | S-H | B-H | P-H |
|---|---|---|---|---|---|---|---|
| **Platyrrhines** | ***Alouatta*** | 0.0117<br>± < 0.0001<br>(0.0116, 0.0118) | 0.0119<br>± < 0.0001<br>(0.0118, 0.0119) | 0.012<br>± < 0.0001<br>(0.0119, 0.012) | 0.0096<br>± < 0.0001<br>(0.0096, 0.0097) | 0.0097<br>± < 0.0001<br>(0.0096, 0.0098) | 0.0099<br>± < 0.0001<br>(0.0098, 0.0100) |
| **Cercopithecoids** | ***Colobus*** | 0.0128<br>± < 0.0001<br>(0.0127, 0.0129) | 0.0129<br>± < 0.0001<br>(0.0128, 0.0129) | 0.0082<br>± < 0.0001<br>(0.0082, 0.0083) | 0.007<br>± < 0.0001<br>(0.0070, 0.0071) | 0.0116<br>± < 0.0001<br>(0.0116, 0.0117) | 0.0073<br>± < 0.0001<br>(0.0072, 0.0073) |
| | ***Macaca*** | 0.0108<br>± < 0.0001<br>(0.0107, 0.0108) | 0.0104<br>± < 0.0001<br>(0.0103, 0.0104) | 0.0135<br>± < 0.0001<br>(0.0134, 0.0135) | 0.0113<br>± < 0.0001<br>(0.0112, 0.0114) | 0.0086<br>± < 0.0001<br>(0.0086, 0.0087) | 0.0117<br>± < 0.0001<br>(0.0117, 0.0118) |
| | ***Papio*** | 0.0257<br>± < 0.0001<br>(0.0256, 0.0259) | 0.0414<br>± < 0.0001<br>(0.0412, 0.0415) | 0.0372<br>± < 0.0001<br>(0.0370, 0.0374) | 0.0379<br>± < 0.0001<br>(0.0378, 0.0381) | 0.0263<br>± < 0.0001<br>(0.0261, 0.0264) | 0.0215<br>± < 0.0001<br>(0.0213, 0.0216) |
| **Hominoids** | ***Homo*** | 0.0095<br>± < 0.0001<br>(0.0095, 0.0096) | 0.0105<br>± < 0.0001<br>(0.0104, 0.0105) | 0.0076<br>± < 0.0001<br>(0.0076, 0.0077) | 0.0065<br>± < 0.0001<br>(0.0065, 0.0065) | 0.0085<br>± < 0.0001<br>(0.0084, 0.0085) | 0.0057<br>± < 0.0001<br>(0.0056, 0.0057) |
| | ***Hylobates*** | 0.0219<br>± < 0.0001<br>(0.0218, 0.0220) | 0.0267<br>± < 0.0001<br>(0.0266, 0.0268) | 0.0157<br>± < 0.0001<br>(0.0156, 0.0158) | 0.0142<br>± < 0.0001<br>(0.0141, 0.0143) | 0.0206<br>± < 0.0001<br>(0.0205, 0.0207) | 0.0094<br>± < 0.0001<br>(0.0093, 0.0095) |
| | ***Pan*** | 0.0096<br>± < 0.0001<br>(0.0095, 0.0097) | 0.0133<br>± < 0.0001<br>(0.0132, 0.0133) | 0.0116<br>± < 0.0001<br>(0.0116, 0.0117) | 0.0106<br>± < 0.0001<br>(0.0105, 0.0107) | 0.0087<br>± < 0.0001<br>(0.0086, 0.0087) | 0.0072<br>± < 0.0001<br>(0.0071, 0.0072) |
| | ***Pongo*** | 0.0351<br>± < 0.0001<br>(0.0350, 0.0353) | 0.0419<br>± < 0.0001<br>(0.0418, 0.0421) | 0.0209<br>± < 0.0001<br>(0.0207, 0.0211) | 0.0206<br>± < 0.0001<br>(0.0204, 0.0207) | 0.0344<br>± < 0.0001<br>(0.0343, 0.0346) | 0.0131<br>± < 0.0001<br>(0.0130, 0.0133) |

Abbreviations: B-P = basicranium and pelvic girdle, B-S = basicranium and shoulder girdle, S-P = shoulder girdle and pelvic girdle, S-H = shoulder girdle and humerus, B-H = basicranium and humerus, P-H = pelvic girdle and humerus.

### SOM Table S7

Mean conditioned covariance, standard error (indicated by ±), and 95% confidence intervals (in parentheses) for paired anatomical regions for each genus using mean standardized variance covariance matrices. Among comparisons of anatomical regions, cells colored light yellow indicate the lower conditioned covariance, whereas cells colored green have a higher conditioned covariance. Uncolored cells indicate comparisons where the conditioned covariances are similar. Lower values reflect smaller responses to directional selection, and thus more constraint being imposed by the set of traits under stabilizing selection. Analyses that deviate from the primary pattern among genera have bolded values.

| **Paired** | **Platyrrhines** | **Cercopithecoids** | | | **Hominoids** | | | |
|---|---|---|---|---|---|---|---|---|
| **regions[a]** | ***Alouatta*** | ***Colobus*** | ***Macaca*** | ***Papio*** | ***Homo*** | ***Hylobates*** | ***Pan*** | ***Pongo*** |
| **B(x), S(y)** | 0.0258 | 0.0287 | **0.0469** | 0.0381 | 0.0371 | 0.0787 | **0.0303** | 0.0653 |
| | ± 0.0003 | ± 0.0003 | **± 0.0002** | ± 0.0007 | ± 0.0002 | ± 0.0004 | **± 0.0003** | ± 0.0006 |
| | (0.0251, 0.0264) | (0.0282, 0.0292) | (0.0465, 0.0472) | (0.0367, 0.0396) | (0.0368, 0.0375) | (0.0780, 0.0794) | (0.0297, 0.0309) | (0.0641, 0.0666) |
| **S(x), B(y)** | 0.0269 | 0.0594 | **0.0288** | 0.0442 | 0.0453 | 0.0896 | **0.0297** | 0.2139 |
| | ± 0.0003 | ± 0.0004 | **± 0.0002** | ± 0.0004 | ± 0.0002 | ± 0.0005 | **± 0.0003** | ± 0.0007 |
| | (0.0269, 0.0269) | (0.0587, 0.0601) | (0.0285, 0.0292) | (0.0434, 0.045) | (0.0450, 0.0457) | (0.0887, 0.0905) | (0.0291, 0.0302) | (0.2126, 0.2152) |
| **B(x), P(y)** | 0.0383 | 0.0353 | **0.0626** | 0.0188 | 0.031 | 0.0501 | 0.0292 | 0.0506 |
| | ± 0.0003 | ± 0.0002 | **± 0.0002** | ± 0.0004 | ± 0.0001 | ± 0.0003 | ± 0.0002 | ± 0.0005 |
| | (0.0383, 0.0383) | (0.0349, 0.0357) | **(0.0622, 0.0631)** | (0.0179, 0.0196) | (0.0308, 0.0313) | (0.0496, 0.0506) | (0.0288, 0.0296) | (0.0496, 0.0516) |
| | 0.0473 | 0.0724 | **0.0362** | 0.0553 | 0.0543 | 0.1449 | 0.0451 | 0.2084 |

| | | | | | | | | |
|---|---|---|---|---|---|---|---|---|
| **P(x), B(y)** | ± 0.0003 (0.0473, 0.0473) | ± 0.0002 (0.0719, 0.0729) | **± 0.0002 (0.0358, 0.0366)** | ± 0.0004 (0.0545, 0.0561) | ± 0.0002 (0.0540, 0.0546) | ± 0.0003 (0.1442, 0.1455) | ± 0.0002 (0.0446, 0.0456) | ± 0.0007 (0.2070, 0.2097) |
| **B(x), H(y)** | 0.0211 ± 0.0003 (0.0211, 0.0211) | 0.0274 ± 0.0002 (0.0270, 0.0278) | **0.0418 ± 0.0002 (0.0414, 0.0421)** | 0.0245 ± 0.0005 (0.0236, 0.0254) | 0.0217 ± 0.0001 (0.0214, 0.0219) | 0.0344 ± 0.0003 (0.0339, 0.0349) | 0.0244 ± 0.0002 (0.0240, 0.0248) | 0.0537 ± 0.0005 (0.0527, 0.0546) |
| **H(x), B(y)** | 0.0415 ± 0.0003 (0.0415, 0.0415) | 0.0758 ± 0.0002 (0.0754, 0.0762) | **0.035 ± 0.0002 (0.0346, 0.0354)** | 0.0584 ± 0.0004 (0.0576, 0.0592) | 0.0568 ± 0.0002 (0.0565, 0.0572) | 0.1457 ± 0.0003 (0.1451, 0.1464) | 0.0464 ± 0.0002 (0.0460, 0.0469) | 0.2219 ± 0.0007 (0.2206, 0.2232) |
| **S(x), H(y)** | 0.0235 ± 0.0002 (0.0235, 0.0235) | 0.0265 ± 0.0002 (0.0261, 0.0269) | 0.0382 ± 0.0002 (0.0379, 0.0385) | 0.0195 ± 0.0004 (0.0188, 0.0202) | 0.0215 ± 0.0001 (0.0213, 0.0217) | 0.033 ± 0.0003 (0.0325, 0.0336) | 0.023 ± 0.0002 (0.0227, 0.0234) | 0.0434 ± 0.0006 (0.0423, 0.0445) |
| **H(x), S(y)** | 0.0335 ± 0.0003 (0.0335, 0.0335) | 0.0347 ± 0.0002 (0.0342, 0.0351) | 0.0532 ± 0.0002 (0.0528, 0.0536) | 0.0273 ± 0.0007 (0.0260, 0.0287) | 0.0403 ± 0.0002 (0.0400, 0.0406) | 0.0833 ± 0.0003 (0.0826, 0.0839) | 0.0541 ± 0.0003 (0.0536, 0.0546) | 0.0982 ± 0.0006 (0.0970, 0.0994) |
| **S(x), P(y)** | **0.0417 ± 0.0002 (0.0417, 0.0417)** | **0.0328 ± 0.0002 (0.0324, 0.0332)** | **0.0515 ± 0.0002 (0.0511, 0.0519)** | 0.0172 ± 0.0004 (0.0165, 0.0180) | 0.0261 ± 0.0001 (0.0258, 0.0263) | 0.0419 ± 0.0003 (0.0414, 0.0425) | 0.0238 ± 0.0002 (0.0234, 0.0242) | 0.0424 ± 0.0005 (0.0413, 0.0434) |
| **P(x), S(y)** | **0.0318 ± 0.0003 (0.0318, 0.0318)** | **0.0278 ± 0.0003 (0.0273, 0.0283)** | **0.0495 ± 0.0002 (0.0491, 0.0499)** | 0.0332 ± 0.0007 (0.0319, 0.0345) | 0.0318 ± 0.0002 (0.0315, 0.0321) | 0.0831 ± 0.0004 (0.0824, 0.0838) | 0.0354 ± 0.0003 (0.0348, 0.0359) | 0.0821 ± 0.0007 (0.0808, 0.0834) |

| | | | | | | | | |
|---|---|---|---|---|---|---|---|---|
| **P(x), H(y)** | 0.0203 ± 0.0003 (0.0203, 0.0203) | 0.0222 ± 0.0002 (0.0218, 0.0227) | 0.0332 ± 0.0002 (0.0329, 0.0336) | **0.0192 ± 0.0005 (0.0182, 0.0202)** | 0.0195 ± 0.0001 (0.0193, 0.0197) | 0.0326 ± 0.0003 (0.0320, 0.0331) | 0.0229 ± 0.0002 (0.0225, 0.0233) | 0.0452 ± 0.0006 (0.0441, 0.0463) |
| **H(x), P(y)** | 0.0242 ± 0.0003 (0.0242, 0.0242) | 0.0336 ± 0.0002 (0.0332, 0.0340) | 0.0483 ± 0.0002 (0.0479, 0.0487) | **0.0158 ± 0.0005 (0.0149, 0.0168)** | 0.0286 ± 0.0001 (0.0283, 0.0288) | 0.0448 ± 0.0003 (0.0443, 0.0453) | 0.0289 ± 0.0002 (0.0285, 0.0294) | 0.049 ± 0.0006 (0.0477, 0.0499) |

Abbreviations: B = basicranium, P = pelvic girdle, S = shoulder girdle, H = humerus.

[a]The anatomical region held constant by stabilizing selection is denoted by (x), and the anatomical region under directional selection is indicated by (y).

**SOM Table S8**

Mean evolutionary flexibility, standard error (indicated by ±), and 95% confidence intervals (in parentheses) for paired anatomical regions among genera using mean standardized variance covariance matrices. Higher values reflect greater flexibility, and thus more evolvability for that grouping of traits compared among genera (rows). Colors indicate the level of evolutionary flexibility compared between genera for each paired anatomical region: lighter shades of green indicate less constraint and darker shades of green indicate more constraint among regions.

| Paired | Platyrrhines | Cercopithecoids | | | Hominoids | | | |
|---|---|---|---|---|---|---|---|---|
| regions[a] | *Alouatta* | *Colobus* | *Macaca* | *Papio* | *Homo* | *Hylobates* | *Pan* | *Pongo* |
| B-P | 0.8092 ± 0.0009 (0.8075, 0.8109) | 0.7895 ± 0.0008 (0.7879, 0.7911) | 0.8450 ± 0.0006 (0.8438, 0.8463) | 0.7589 ± 0.0005 (0.7580, 0.7599) | 0.8241 ± 0.0003 (0.8234, 0.8247)) | 0.7563 ± 0.0007 (0.7549, 0.7576) | 0.8084 ± 0.0007 (0.8070, 0.8099) | 0.7156 ± 0.0008 (0.7141, 0.7172) |
| B-S | 0.7736 ± 0.0008 (0.7720, 0.7753) | 0.7817 ± 0.0008 (0.7800, 0.7833) | 0.8134 ± 0.0006 (0.8121, 0.8146) | 0.7488 ± 0.0004 (0.7480, 0.7497) | 0.6679 ± 0.0005 (0.6669, 0.6689) | 0.7779 ± 0.0005 (0.7770, 0.7789) | 0.7516 ± 0.0007 (0.7503, 0.7530) | 0.7511 ± 0.0006 (0.7499, 0.7523) |
| S-P | 0.7928 ± 0.0008 (0.7912, 0.7944) | 0.8462 ± 0.0009 (0.8444, 0.8480) | 0.8223 ± 0.0006 (0.8211, 0.8234) | 0.7348 ± 0.0006 (0.7337, 0.7359) | 0.7034 ± 0.0005 (0.7025, 0.7043) | 0.7942 ± 0.0009 (0.7924, 0.7960) | 0.7670 ± 0.0008 (0.7655, 0.7685) | 0.7625 ± 0.0010 (0.7605, 0.7646) |
| S-H | 0.7892 ± 0.0009 (0.7874, 0.7910) | 0.8580 ± 0.0011 (0.8559, 0.8601) | 0.8313 ± 0.0006 (0.8301, 0.8326) | 0.7215 ± 0.0005 (0.7205, 0.7225) | 0.6516 ± 0.0002 (0.6512, 0.652) | 0.7747 ± 0.0010 (0.7728, 0.7767) | 0.7637 ± 0.0009 (0.7619, 0.7655) | 0.769 ± 0.0012 (0.7667, 0.7713) |

| | | | | | | | | |
|---|---|---|---|---|---|---|---|---|
| **B-H** | 0.7927 ± 0.0010 (0.7907, 0.7946) | 0.7659 ± 0.0009 (0.7640, 0.7677) | 0.8673 ± 0.0008 (0.8658, 0.8688) | 0.7490 ± 0.0005 (0.7480, 0.7500) | 0.566 ± 0.0004 (0.5653, 0.5668) | 0.7245 ± 0.0008 (0.7231, 0.7260) | 0.7936 ± 0.0009 (0.7919, 0.7953) | 0.7107 ± 0.0009 (0.7089, 0.7124) |
| **P-H** | 0.7997 ± 0.0009 (0.7979, 0.8016) | 0.8533 ± 0.0010 (0.8513, 0.8552) | 0.8354 ± 0.0007 (0.8341, 0.8367) | 0.7360 ± 0.0006 (0.7348, 0.7371) | 0.6035 ± 0.0004 (0.6027, 0.6043) | 0.8796 ± 0.0009 (0.8778, 0.8813) | 0.8696 ± 0.0008 (0.8680, 0.8712) | 0.8547 ± 0.0011 (0.8525, 0.8569) |
| **All** | 0.7320 ± 0.0009 (0.7303, 0.7337) | 0.7453 ± 0.0010 (0.7434, 0.7472) | 0.7646 ± 0.0006 (0.7634, 0.7658) | 0.7037 ± 0.0004 (0.7030, 0.7044) | 0.6082 ± 0.0003 (0.6076, 0.6089) | 0.7074 ± 0.0007 (0.7060, 0.7088) | 0.7256 ± 0.0008 (0.7240, 0.7271) | 0.6814 ± 0.0009 (0.6797, 0.6831) |

[a]B-PG = basicranium and pelvic girdle, B-S = basicranium and shoulder girdle, S-P = shoulder girdle and pelvic girdle, S-H = shoulder girdle and humerus, B-H = basicranium and humerus, P-H = pelvic girdle and humerus, All = all four regions together.

**SOM Table S9**

Significance for measures of mean evolutionary flexibility (EF), mean variance of eigenvalues (VE), and mean trait autonomy (TA), using size corrected variance covariance matrices. Comparisons are between taxa for each trait grouping: basicranium and pelvic girdle, basicranium and shoulder girdle, shoulder girdle and pelvic girdle, basicranium and humerus, pelvic girdle and humerus, and shoulder girdle and humerus. Comparisons of mean evolutionary flexibility (EF) are in the lower triangle of the upper matrix, comparisons of mean variance of eigenvalues (VE) are in the lower triangle of the lower matrix, and comparisons of mean trait autonomy (TA) are in the upper triangle of the lower matrix. Significant comparisons ($p < 0.05$) are highlighted blue.

### A. Basicranium–Pelvic girdle

| **EF** | *Alouatta* | *Colobus* | *Homo* | *Hylobates* | *Macaca* | *Pan* | *Papio* | *Pongo* |
|---|---|---|---|---|---|---|---|---|
| *Alouatta* | | | | | | | | |
| *Colobus* | 0.098 | | | | | | | |
| *Homo* | 0 | 0.021 | | | | | | |
| *Hylobates* | 0.041 | 0.655 | 0.016 | | | | | |
| *Macaca* | 0.120 | 0.237 | 0.001 | 0.100 | | | | |
| *Pan* | 0.355 | 0.02 | 0 | 0.008 | 0.031 | | | |
| *Papio* | 0.343 | 0.018 | 0 | 0.008 | 0.022 | 0.062 | | |
| *Pongo* | 0.375 | 0.457 | 0.006 | 0.225 | 0.593 | 0.138 | 0.13 | |

| | *Alouatta* | *Colobus* | *Homo* | *Hylobates* | *Macaca* | *Pan* | *Papio* | *Pongo* | **TA** |
|---|---|---|---|---|---|---|---|---|---|
| *Alouatta* | | 0.375 | 0 | 0.068 | 0.283 | 0.029 | 0.378 | 0.759 | *Alouatta* |
| *Colobus* | 0.086 | | 0.019 | 0.795 | 0.723 | 0.451 | 0.276 | 0.591 | *Colobus* |
| *Homo* | 0 | 0.001 | | 0.012 | 0.282 | 0 | 0 | 0.001 | *Homo* |
| *Hylobates* | 0.044 | 0.134 | 0.001 | | 0.688 | 0.118 | 0.024 | 0.239 | *Hylobates* |
| *Macaca* | 0.139 | 0.306 | 0.001 | 0.183 | | 0.333 | 0.221 | 0.472 | *Macaca* |
| *Pan* | 0.332 | 0.323 | 0 | 0.181 | 0.494 | | 0.001 | 0.648 | *Pan* |
| *Papio* | 0.573 | 0.042 | 0 | 0.018 | 0.093 | 0.352 | | 0.151 | *Papio* |
| *Pongo* | 0.356 | 0.042 | 0 | 0.021 | 0.073 | 0.340 | 0.379 | | *Pongo* |
| **VE** | *Alouatta* | *Colobus* | *Homo* | *Hylobates* | *Macaca* | *Pan* | *Papio* | *Pongo* | |

### B. Basicranium–Shoulder girdle

| **EF** | *Alouatta* | *Colobus* | *Homo* | *Hylobates* | *Macaca* | *Pan* | *Papio* | *Pongo* |
|---|---|---|---|---|---|---|---|---|
| *Alouatta* | | | | | | | | |
| *Colobus* | 0.147 | | | | | | | |
| *Homo* | 0.002 | 0.032 | | | | | | |
| *Hylobates* | 0.012 | 0.158 | 0.916 | | | | | |
| *Macaca* | 0.004 | 0.053 | 0.597 | 0.228 | | | | |
| *Pan* | 0.007 | 0.048 | 0.496 | 0.231 | 0.545 | | | |
| *Papio* | 0.089 | 0.362 | 0.059 | 0.353 | 0.119 | 0.124 | | |
| *Pongo* | 0.041 | 0.226 | 0.104 | 0.532 | 0.189 | 0.204 | 0.404 | |

| | *Alouatta* | *Colobus* | *Homo* | *Hylobates* | *Macaca* | *Pan* | *Papio* | *Pongo* | **TA** |
|---|---|---|---|---|---|---|---|---|---|
| *Alouatta* | | 0.643 | 0.141 | 0.692 | 0.394 | 0.778 | 0.517 | 0.848 | *Alouatta* |
| *Colobus* | 0.340 | | 0.007 | 0.242 | 0.282 | 0.013 | 0.485 | 0.924 | *Colobus* |
| *Homo* | 0.025 | 0 | | 0.022 | 0.685 | 0.082 | 0 | 0.193 | *Homo* |
| *Hylobates* | 0.489 | 0.318 | 0.021 | | 0.299 | 0.027 | 0.091 | 0.579 | *Hylobates* |
| *Macaca* | 0.028 | 0 | 0.212 | 0.023 | | 0.364 | 0.252 | 0.436 | *Macaca* |
| *Pan* | 0.196 | 0.036 | 0.269 | 0.302 | 0.279 | | 0.001 | 0.758 | *Pan* |
| *Papio* | 0.303 | 0.11 | 0.001 | 0.516 | 0.002 | 0.177 | | 0.270 | *Papio* |
| *Pongo* | 0.316 | 0.119 | 0.002 | 0.546 | 0.004 | 0.170 | 0.347 | | *Pongo* |
| **VE** | *Alouatta* | *Colobus* | *Homo* | *Hylobates* | *Macaca* | *Pan* | *Papio* | *Pongo* | |

### c. Shoulder girdle–Pelvic girdle

| **EF** | *Alouatta* | *Colobus* | *Homo* | *Hylobates* | *Macaca* | *Pan* | *Papio* | *Pongo* |
|---|---|---|---|---|---|---|---|---|
| *Alouatta* | | | | | | | | |
| *Colobus* | 0.519 | | | | | | | |
| *Homo* | 0.173 | 0.210 | | | | | | |
| *Hylobates* | 0.358 | 0.402 | 0.435 | | | | | |
| *Macaca* | 0.174 | 0.202 | 0.249 | 0.480 | | | | |
| *Pan* | 0.206 | 0.247 | 0.252 | 0.483 | 0.299 | | | |
| *Papio* | 0.365 | 0.211 | 0.054 | 0.100 | 0.044 | 0.048 | | |
| *Pongo* | 0.054 | 0.070 | 0.195 | 0.135 | 0.075 | 0.076 | 0.021 | |

| | *Alouatta* | *Colobus* | *Homo* | *Hylobates* | *Macaca* | *Pan* | *Papio* | *Pongo* | **TA** |
|---|---|---|---|---|---|---|---|---|---|
| *Alouatta* | | 0.523 | 0 | 0.118 | 0.276 | 0.016 | 0.459 | 0.998 | *Alouatta* |
| *Colobus* | 0.129 | | 0.003 | 0.654 | 0.355 | 0.089 | 0.291 | 0.998 | *Colobus* |
| *Homo* | 0.006 | 0 | | 0 | 0.415 | 0.011 | 0 | 0.612 | *Homo* |
| *Hylobates* | 0.111 | 0.316 | 0.012 | | 0.312 | 0.047 | 0.021 | 0.072 | *Hylobates* |
| *Macaca* | 0.023 | 0.006 | 0.074 | 0.081 | | 0.436 | 0.223 | 0.584 | *Macaca* |
| *Pan* | 0.046 | 0.075 | 0.079 | 0.322 | 0.233 | | 0 | 0.143 | *Pan* |
| *Papio* | 0.582 | 0.136 | 0 | 0.114 | 0.003 | 0.998 | | 0.001 | *Papio* |
| *Pongo* | 0 | 0 | 0.037 | 0 | 0.003 | 0 | 0 | | *Pongo* |
| **VE** | *Alouatta* | *Colobus* | *Homo* | *Hylobates* | *Macaca* | *Pan* | *Papio* | *Pongo* | |

#### D. Basicranium–Humerus

| EF | *Alouatta* | *Colobus* | *Homo* | *Hylobates* | *Macaca* | *Pan* | *Papio* | *Pongo* |
|---|---|---|---|---|---|---|---|---|
| *Alouatta* | | | | | | | | |
| *Colobus* | 0.810 | | | | | | | |
| *Homo* | 0.104 | 0.348 | | | | | | |
| *Hylobates* | 0.345 | 0.922 | 0.233 | | | | | |
| *Macaca* | 0.687 | 0.878 | 0.593 | 0.818 | | | | |
| *Pan* | 0.892 | 0.662 | 0.056 | 0.204 | 0.565 | | | |
| *Papio* | 0.801 | 0.364 | 0.028 | 0.072 | 0.336 | 0.024 | | |
| *Pongo* | 0.567 | 0.051 | 0.001 | 0.012 | 0.091 | 0 | 0.786 | |

| | *Alouatta* | *Colobus* | *Homo* | *Hylobates* | *Macaca* | *Pan* | *Papio* | *Pongo* | TA |
|---|---|---|---|---|---|---|---|---|---|
| *Alouatta* | | 0.806 | 0.037 | 0.513 | 0.650 | 0.732 | 0.313 | 0.609 | *Alouatta* |
| *Colobus* | 0.039 | | 0.147 | 0.869 | 0.803 | 0.678 | 0.241 | 0.500 | *Colobus* |
| *Homo* | 0.004 | 0.002 | | 0.089 | 0.972 | 0.012 | 0 | 0.001 | *Homo* |
| *Hylobates* | 0.031 | 0.032 | 0.042 | | 0.754 | 0.322 | 0.024 | 0.112 | *Hylobates* |
| *Macaca* | 0.008 | 0.005 | 0.012 | 0.062 | | 0.527 | 0.206 | 0.352 | *Macaca* |
| *Pan* | 0.702 | 0.614 | 0.162 | 0.561 | 0.236 | | 0 | 0.02 | *Pan* |
| *Papio* | 0.001 | 0 | 0 | 0 | 0 | 0.002 | | 0.134 | *Papio* |
| *Pongo* | 0 | 0 | 0 | 0 | 0 | 0.003 | 0.998 | | *Pongo* |
| VE | *Alouatta* | *Colobus* | *Homo* | *Hylobates* | *Macaca* | *Pan* | *Papio* | *Pongo* | |

### E. Pelvic girdle–Humerus

| EF | *Alouatta* | *Colobus* | *Homo* | *Hylobates* | *Macaca* | *Pan* | *Papio* | *Pongo* |
|---|---|---|---|---|---|---|---|---|
| *Alouatta* | | | | | | | | |
| *Colobus* | 0.237 | | | | | | | |
| *Homo* | 0.002 | 0.018 | | | | | | |
| *Hylobates* | 0.238 | 0.415 | 0.003 | | | | | |
| *Macaca* | 0.703 | 0.019 | 0 | 0.050 | | | | |
| *Pan* | 0.700 | 0.019 | 0 | 0.050 | 0.094 | | | |
| *Papio* | 0.630 | 0.010 | 0 | 0.025 | 0.117 | 0.051 | | |
| *Pongo* | 0.175 | 0.471 | 0.115 | 0.272 | 0.049 | 0.049 | 0.034 | |

| | *Alouatta* | *Colobus* | *Homo* | *Hylobates* | *Macaca* | *Pan* | *Papio* | *Pongo* | **TA** |
|---|---|---|---|---|---|---|---|---|---|
| *Alouatta* | | 0.417 | 0 | 0.072 | 0.359 | 0.034 | 0.430 | 0.990 | *Alouatta* |
| *Colobus* | 0.029 | | 0.008 | 0.848 | 0.897 | 0.611 | 0.291 | 0.967 | *Colobus* |
| *Homo* | 0 | 0 | | 0.008 | 0.025 | 0 | 0 | 0.185 | *Homo* |
| *Hylobates* | 0.013 | 0.223 | 0.004 | | 0.897 | 0.212 | 0.028 | 0.427 | *Hylobates* |
| *Macaca* | 0.001 | 0.016 | 0.007 | 0.176 | | 0.637 | 0.248 | 0.543 | *Macaca* |
| *Pan* | 0.002 | 0.049 | 0.041 | 0.333 | 0.459 | | 0 | 0.185 | *Pan* |
| *Papio* | 0.418 | 0.001 | 0 | 0 | 0 | 1 | | 0.001 | *Papio* |
| *Pongo* | 0 | 0 | 1 | 0 | 0 | 0.001 | 0 | | *Pongo* |
| **VE** | *Alouatta* | *Colobus* | *Homo* | *Hylobates* | *Macaca* | *Pan* | *Papio* | *Pongo* | |

### F. Shoulder girdle–Humerus

| **EF** | *Alouatta* | *Colobus* | *Homo* | *Hylobates* | *Macaca* | *Pan* | *Papio* | *Pongo* |
|---|---|---|---|---|---|---|---|---|
| *Alouatta* | | | | | | | | |
| *Colobus* | 0.220 | | | | | | | |
| *Homo* | 0.063 | 0.227 | | | | | | |
| *Hylobates* | 0.208 | 0.446 | 0.167 | | | | | |
| *Macaca* | 0.046 | 0.156 | 0.520 | 0.116 | | | | |
| *Pan* | 0.018 | 0.067 | 0.249 | 0.044 | 0.438 | | | |
| *Papio* | 0.742 | 0.924 | 0.406 | 0.873 | 0.320 | 0.129 | | |
| *Pongo* | 0.014 | 0.054 | 0.211 | 0.036 | 0.395 | 0.456 | 0.106 | |

| | *Alouatta* | *Colobus* | *Homo* | *Hylobates* | *Macaca* | *Pan* | *Papio* | *Pongo* | **TA** |
|---|---|---|---|---|---|---|---|---|---|
| *Alouatta* | | 0.638 | 0 | 0.146 | 0.291 | 0.010 | 0.455 | 1 | *Alouatta* |
| *Colobus* | 0.394 | | 0.001 | 0.167 | 0.299 | 0.014 | 0.290 | 1 | *Colobus* |
| *Homo* | 0.026 | 0.006 | | 0.006 | 0.798 | 0.095 | 0 | 0.466 | *Homo* |
| *Hylobates* | 0.769 | 0.518 | 0.069 | | 0.337 | 0.021 | 0.029 | 0.118 | *Hylobates* |
| *Macaca* | 0.020 | 0.003 | 0.359 | 0.054 | | 0.578 | 0.227 | 0.503 | *Macaca* |
| *Pan* | 0.024 | 0.006 | 0.406 | 0.066 | 0.548 | | 0 | 0.283 | *Pan* |
| *Papio* | 0.166 | 0.406 | 0 | 0.780 | 0 | 0.972 | | 0 | *Papio* |
| *Pongo* | 0 | 0 | 0.007 | 0 | 0.005 | 0.054 | 0 | | *Pongo* |
| **VE** | *Alouatta* | *Colobus* | *Homo* | *Hylobates* | *Macaca* | *Pan* | *Papio* | *Pongo* | |

**SOM Table S10.**

Mean trait autonomy (TA), standard error (indicated by ±), and 95% confidence intervals (in parentheses) by taxa for pairwise trait groupings among anatomical regions (columns) using size corrected variance covariance matrices. Larger TA values reflect greater autonomy, and thus less integration, among the traits. Comparisons among traits within taxa are in rows. Colors reflect the degree of integration, and thus constraint, between pairs of anatomical regions within each taxon: lighter shades indicate greater autonomy among traits and darker shades indicate lower autonomy among traits.

| | Taxon | B-P | B-S | S-P | S-H | B-H | P-H | All |
|---|---|---|---|---|---|---|---|---|
| **Platyrrhines** | ***Alouatta*** | 0.0186 ± 0.0005 (0.0176, 0.0196) | 0.0262 ± 0.0004 (0.0254, 0.0270) | 0.0121 ± 0.0003 (0.0115, 0.0126) | 0.0172 ± 0.0003 (0.0165, 0.0179) | 0.0504 ± 0.0007 (0.0490, 0.0517) | 0.0163 ± 0.0004 (0.0155, 0.0171) | 0.0056 ± 0.0004 (0.0048, 0.0065) |
| **Cercopithecoids** | ***Colobus*** | 0.0418 ± 0.0006 (0.0406, 0.0429) | 0.0167 ± 0.0003 (0.0161, 0.0174) | 0.0171 ± 0.0003 (0.0165, 0.0176) | 0.0181 ± 0.0003 (0.0174, 0.0188) | 0.0598 ± 0.0007 (0.0585, 0.0611) | 0.0388 ± 0.0005 (0.0378, 0.0399) | 0.0097 ± 0.0003 (0.0092, 0.0102) |
| | ***Macaca*** | 0.0618 ± 0.0006 (0.0605, 0.063) | 0.0442 ± 0.0006 (0.0430, 0.0454) | 0.0401 ± 0.0005 (0.0392, 0.0411) | 0.044 ± 0.0006 (0.0429, 0.0451) | 0.0795 ± 0.0007 (0.0780, 0.0809) | 0.0437 ± 0.0005 (0.0428, 0.0447) | 0.0186 ± 0.0005 (0.0177, 0.0195) |
| | ***Papio*** | 0.0104 ± 0.0003 (0.0097, 0.0110) | 0.0118 ± 0.0002 (0.0114, 0.0122) | 0.0055 ± 0.0002 (0.0050, 0.0059) | 0.0068 ± 0.0002 (0.0064, 0.0071) | 0.0105 ± 0.0003 (0.0099, 0.0111) | 0.0053 ± 0.0003 (0.0047, 0.0059) | 0.0019 ± 0.0003 (0.0014, 0.0024) |
| **Hominoids** | ***Homo*** | 0.0793 ± 0.0004 (0.0785, 0.0801) | 0.0393 ± 0.0004 (0.0385, 0.0401) | 0.0412 ± 0.0004 (0.0404, 0.0421) | 0.0421 ± 0.0004 (0.0413, 0.0430) | 0.0787 ± 0.0004 (0.0779, 0.0794) | 0.0776 ± 0.0004 (0.0768, 0.0785) | 0.0231 ± 0.0002 (0.0227, 0.0235) |
| | ***Hylobates*** | 0.0401 | 0.0190 | 0.0149 | 0.0204 | 0.0562 | 0.0385 | 0.0079 |

| | | | | | | | |
|---|---|---|---|---|---|---|---|
| | ± 0.0005 | ± 0.0003 | ± 0.0003 | ± 0.0004 | ± 0.0005 | ± 0.0005 | ± 0.0001 |
| | (0.0392, 0.0410) | (0.0184, 0.0197) | (0.0143, 0.0155) | (0.0197, 0.0212) | (0.0552, 0.0572) | (0.0375, 0.0394) | (0.0077, 0.0082) |
| | 0.0227 | 0.0236 | 0.0198 | 0.0310 | 0.0434 | 0.0255 | 0.0071 |
| ***Pan*** | ± 0.0005 | ± 0.0005 | ± 0.0003 | ± 0.0005 | ± 0.0005 | ± 0.0005 | ± 0.0002 |
| | (0.0218, 0.0237) | (0.0227, 0.0246) | (0.0191, 0.0205) | (0.0301, 0.0319) | (0.0424, 0.0443) | (0.0246, 0.0264) | (0.0068, 0.0075) |
| | 0.0290 | 0.0281 | 0.0511 | 0.0701 | 0.0330 | 0.0560 | 0.0149 |
| ***Pongo*** | ± 0.0003 | ± 0.0003 | ± 0.0005 | ± 0.0006 | ± 0.0003 | ± 0.0006 | ± 0.0002 |
| | (0.0284, 0.0297) | (0.0275, 0.0287) | (0.0501, 0.0521) | (0.0690, 0.0712) | (0.0324, 0.0337) | (0.0549, 0.0570) | (0.0146, 0.0152) |

Abbreviations: B-P = Basicranium and pelvic girdle, B-S = basicranium and shoulder girdle, S-P = shoulder girdle and pelvic girdle, S-H = shoulder girdle and humerus, B-H = basicranium and humerus, P-H = pelvic girdle and humerus, All = all four regions.

**SOM Table S11**

Mean variance of the eigenvalues (VE), standard errors (indicated by ±), and 95% confidence intervals (in parentheses) by taxa for pairwise trait groupings among anatomical regions (columns) using mean standardized variance covariance matrices. Greater VE values reflect more integration among the traits. Comparisons among traits within genera are in rows. Colors reflect the degree of integration, and thus constraint, between pairs of anatomical regions within each taxon: lighter shades of green indicate weaker integration among traits and darker shades of green indicate stronger integration among traits.

| | Taxon | B-P | B-S | S-P | S-H | B-H | P-H | All |
|---|---|---|---|---|---|---|---|---|
| **Platyrrhines** | ***Alouatta*** | 0.0096 ± < 0.0001 (0.0095, 0.0098) | 0.0176 ± < 0.0001 (0.0174, 0.0178) | 0.0158 ± < 0.0001 (0.0156, 0.0159) | 0.0168 ± 0.0001 (0.0166, 0.017) | 0.0122 ± < 0.0001 (0.0121, 0.0124) | 0.0097 ± < 0.0001 (0.0096, 0.0098) | 0.0062 ± < 0.0001 (0.0062, 0.0063) |
| **Cercopithecoids** | ***Colobus*** | 0.0133 ± < 0.0001 (0.0131, 0.0134) | 0.0149 ± < 0.0001 (0.0148, 0.0151) | 0.0091 ± < 0.0001 (0.009, 0.0093) | 0.0069 ± < 0.0001 (0.0068, 0.007) | 0.0162 ± < 0.0001 (0.016, 0.0164) | 0.0048 ± < 0.0001 (0.0047, 0.0049) | 0.0043 ± < 0.0001 (0.0043, 0.0044) |
| | ***Macaca*** | 0.0074 ± < 0.0001 (0.0073, 0.0075) | 0.0153 ± < 0.0001 (0.0151, 0.0155) | 0.0099 ± < 0.0001 (0.0098, 0.0101) | 0.0109 ± < 0.0001 (0.0107, 0.011) | 0.0074 ± < 0.0001 (0.0073, 0.0075) | 0.0065 ± < 0.0001 (0.0064, 0.0066) | 0.0045 ± < 0.0001 (0.0045, 0.0046) |
| | ***Papio*** | 0.0361 ± 0.0001 (0.0359, 0.0364) | 0.0389 ± 0.0001 (0.0387, 0.0391) | 0.0501 ± 0.0001 (0.0498, 0.0503) | 0.0449 ± 0.0001 (0.0446, 0.0452) | 0.0276 ± 0.0001 (0.0274, 0.0278) | 0.0308 ± < 0.0001 (0.0306, 0.031) | 0.0204 ± < 0.0001 (0.0203, 0.0205) |
| **Hominoids** | ***Homo*** | 0.0077 | 0.0379 | 0.038 | 0.0506 | 0.0493 | 0.0277 | 0.0174 |

| | | | | | | | |
|---|---|---|---|---|---|---|---|
| | ± < 0.0001 (0.0077, 0.0078) | ± 0.0001 (0.0376, 0.0381) | ± 0.0001 (0.0378, 0.0383) | ± 0.0001 (0.0504, 0.0508) | ± 0.0001 (0.0491, 0.0495) | ± < 0.0001 (0.0276, 0.0278) | ± < 0.0001 (0.0173, 0.0175) |
| ***Hylobates*** | 0.0213 ± 0.0001 (0.0211, 0.0215) | 0.0196 ± < 0.0001 (0.0194, 0.0198) | 0.0135 ± < 0.0001 (0.0134, 0.0137) | 0.0156 ± 0.0001 (0.0154, 0.0158) | 0.025 ± 0.0001 (0.0248, 0.0253) | 0.0071 ± < 0.0001 (0.007, 0.0072) | 0.0064 ± < 0.0001 (0.0064, 0.0065) |
| ***Pan*** | 0.0133 ± < 0.0001 (0.0131, 0.0134) | 0.0193 ± < 0.0001 (0.0192, 0.0195) | 0.0163 ± < 0.0001 (0.0162, 0.0165) | 0.015 ± < 0.0001 (0.0149, 0.0151) | 0.0156 ± < 0.0001 (0.0154, 0.0158) | 0.0083 ± < 0.0001 (0.0083, 0.0084) | 0.0061 ± < 0.0001 (0.006, 0.0061) |
| ***Pongo*** | 0.0214 ± < 0.0001 (0.0212, 0.0216) | 0.017 ± < 0.0001 (0.0169, 0.0172) | 0.0163 ± 0.0001 (0.0161, 0.0165) | 0.0145 ± 0.0001 (0.0143, 0.0147) | 0.0216 ± 0.0001 (0.0214, 0.0218) | 0.0085 ± < 0.0001 (0.0084, 0.0086) | 0.0068 ± < 0.0001 (0.0067, 0.0068) |

Abbreviations: B-P = Basicranium and pelvic girdle, B-S = basicranium and shoulder girdle, S-P = shoulder girdle and pelvic girdle, S-H = shoulder girdle and humerus, B-H = basicranium and humerus, P-H = pelvic girdle and humerus, All = all four regions.

**SOM Table S12**

Mean trait autonomy (TA), standard error (indicated by ±), and 95% confidence intervals (in parentheses) by genus for pairwise trait groupings among anatomical regions (columns) using mean standardized variance covariance matrices. Greater TA values reflect greater autonomy, and thus less integration, among the traits. Colors reflect the degree of integration, and thus constraint, between pairs of anatomical regions within each taxon: lighter shades indicate greater autonomy among traits and darker shades indicate lower autonomy among traits.

| | Taxon | B–P | B–S | S–P | S–H | B–H | P–H | All |
|---|---|---|---|---|---|---|---|---|
| **Platyrrhines** | ***Alouatta*** | 0.0077<br>± 0.0002<br>(0.0072, 0.0081) | 0.0220<br>± 0.0004<br>(0.0211, 0.0228) | 0.0337<br>± 0.0004<br>(0.0329, 0.0346) | 0.0253<br>± 0.0004<br>(0.0245, 0.0261) | 0.0331<br>± 0.0004<br>(0.0323, 0.0339) | 0.0270<br>± 0.0004<br>(0.0262, 0.0279) | 0.0285<br>± 0.0004<br>(0.0277, 0.0293) |
| **Cercopithecoids** | ***Colobus*** | 0.0167<br>± 0.0002<br>(0.0163, 0.0172) | 0.0333<br>± 0.0004<br>(0.0325, 0.0341) | 0.0371<br>± 0.0004<br>(0.0363, 0.0378) | 0.0511<br>± 0.0004<br>(0.0503, 0.0519) | 0.0441<br>± 0.0004<br>(0.0433, 0.0448) | 0.0533<br>± 0.0004<br>(0.0525, 0.0541) | 0.0605<br>± 0.0004<br>(0.0597, 0.0612) |
| | ***Macaca*** | 0.0250<br>± 0.0001<br>(0.0247, 0.0252) | 0.0568<br>± 0.0003<br>(0.0563, 0.0574) | 0.0730<br>± 0.0003<br>(0.0724, 0.0737) | 0.0554<br>± 0.0003<br>(0.0548, 0.0559) | 0.0736<br>± 0.0003<br>(0.0729, 0.0742) | 0.0617<br>± 0.0003<br>(0.0612, 0.0623) | 0.0593<br>± 0.0002<br>(0.0588, 0.0598) |
| | ***Papio*** | 0.0019<br>± < 0.0001<br>(0.0018, 0.0021) | 0.0041<br>± 0.0002<br>(0.0037, 0.0045) | 0.0084<br>± 0.0003<br>(0.0079, 0.0090) | 0.0045<br>± 0.0002<br>(0.0042, 0.0049) | 0.0096<br>± 0.0003<br>(0.0090, 0.0102) | 0.0109<br>± 0.0003<br>(0.0104, 0.0114) | 0.0043<br>± 0.0002<br>(0.0040, 0.0047) |
| **Hominoids** | ***Homo*** | 0.0151<br>± < 0.0001<br>(0.0150, 0.0152) | 0.0381<br>± 0.0001<br>(0.0378, 0.0383) | 0.0594<br>± 0.0002<br>(0.0591, 0.0597) | 0.0354<br>± 0.0001<br>(0.0352, 0.0356) | 0.0446<br>± 0.0001<br>(0.0443, 0.0448) | 0.0361<br>± < 0.0001<br>(0.0359, 0.0363) | 0.0326<br>± < 0.0001<br>(0.0324, 0.0327) |
| | ***Hylobates*** | 0.0156 | 0.0281 | 0.0329 | 0.0563 | 0.0301 | 0.0722 | 0.0632 |

| | | | | | | | |
|---|---|---|---|---|---|---|---|
| | ± 0.0001 (0.0153, 0.0159) | ± 0.0002 (0.0277, 0.0286) | ± 0.0003 (0.0324, 0.0334) | ± 0.0003 (0.0557, 0.0569) | ± 0.0003 (0.0295, 0.0306) | ± 0.0003 (0.0716, 0.0727) | ± 0.0003 (0.0626, 0.0637) |
| ***Pan*** | 0.0127 ± 0.0002 (0.0123, 0.0131) | 0.0241 ± 0.0003 (0.0234, 0.0247) | 0.0386 ± 0.0003 (0.0379, 0.0393) | 0.0366 ± 0.0003 (0.036, 0.0372) | 0.0457 ± 0.0003 (0.045, 0.0464) | 0.0606 ± 0.0003 (0.06, 0.0612) | 0.0484 ± 0.0003 (0.0478, 0.049) |
| ***Pongo*** | 0.0130 ± 0.0002 (0.0126, 0.0133) | 0.0241 ± 0.0003 (0.0234, 0.0248) | 0.0338 ± 0.0003 (0.0331, 0.0345) | 0.0418 ± 0.0004 (0.0409, 0.0427) | 0.0362 ± 0.0004 (0.0354, 0.0369) | 0.0643 ± 0.0004 (0.0636, 0.0651) | 0.0535 ± 0.0004 (0.0527, 0.0543) |

Abbreviations: B-P = Basicranium and pelvic girdle, B-S = basicranium and shoulder girdle, S-P = shoulder girdle and pelvic girdle, S-H = shoulder girdle and humerus, B-H = basicranium and humerus, P-H = pelvic girdle and humerus, All = all four regions.

**SOM References**

Agosto, E.R., Auerbach, B.M., 2021. Evolvability and constraint in the primate basicranium, shoulder, and hip and the importance of multi-trait evolution. Evol. Biol. 48, 221-232. doi: 10.1007/s11692-021-09532-2.

Berge, C., Kazmierczak, J.B., 1986. Effects of size and locomotor adaptations on the hominid pelvis: evaluation of australopithecine bipedality with a new multivariate method. Folia Primatol. 46,185–204.

Bramble, D.M., Lieberman, D.E., 2004. Endurance running and the evolution of *Homo*. Nature 432, 345–352.

de Oliveira, F.B., Porto, A., Marroig, G., 2009. Covariance structure in the skull of Catarrhini: A case of pattern stasis and magnitude evolution. J. Hum. Evol. 56, 417–430.

Diogo, R., Wood, B.A., 2012. Comparative anatomy and phylogeny of primate muscles and human evolution. CRC Press, Boca Raton.

Grabowski, M., Porto, G., 2017. How many more? Sample size determination in studies of morphological integration and evolvability. Methods Ecol. Evol. 8, 592–603.

Grabowski, M., Roseman, C.C., 2015. Complex and changing patterns of natural selection explain the evolution of the human hip. J. Hum. Evol. 85, 94–110.

Huang, R., Zhi, Q., Patel, K., Wilting, J., Christ, B., 2000. Dual origin and segmental organization of the avian scapula. Development 127, 3789–3794.

Huang, R., Christ, B., Patel, K., 2006. Regulation of scapula development. Anat. Embryol. 211, S65–S71.

Lewton, K.L., 2012. Evolvability of the primate pelvic girdle. Evol. Biol. 39, 126–139.

Little, R.J.A., Rubin, D.B., 2019. Statistical analysis with missing data. Third Edition. John Wiley & Sons, Hoboken.

Lovejoy, C.O., 2005. The natural history of human gait and posture. Part 1. Spine and pelvis. Gait Posture 21, 95–112.

Lovejoy, C.O., Suwa, G., Spurlock, L., Asfaw, B., White, T.D., 2009. The pelvis and femur of *Australopithecus ramidus*: The emergence of upright walking. Science 326, 71e1–71e6.

Marroig, G., Melo, D.A.R., Garcia, G., 2012. Modularity, noise, and natural selection. Evolution 66, 1506–1524.

Martin, R., 1928. Lehrbuch der anthropologie in systematischer darstellung mit besonderer berücksichtigung der anthropologischen methoden für studierende ärtze und forschungsreisende. Zweiter Band: Kraniologie, Osteologie, 2nd ed. Gustav Fischer, Jena.

Matsuoka, T., Ahlberg, P.E., Kessaris, N., Iannarelli, P., Dennehy, U., Richardson, W.D., McMahon, A.P., Koentges, G., 2005. Neural crest origins of the neck and shoulder. Nature 436, 347–355.

McCollum, M.A., Rosenman, B.A., Suwa, G., Meindl, R.S., Lovejoy, C.O., 2010. The vertebral formula of the last common ancestor of African apes and humans. J. Exp. Zool. 314B, 123–134.

Moore-Jansen, P.M., Ousley, S.D., Jantz, R.L., 1994. Data collection procedures for forensic skeletal material: report of investigations No. 48. Forensic Anthropology Center, Department of Anthropology, The University of Tennessee, Knoxville.

Pilbeam, D., 2004. The Anthropoid postcranial axial skeleton: Comments on development, variation, and evolution. J. Exp. Zool. 302B, 241–267.

Richmond, F.J.R., Singh, K., Corneil, B.D., 2001. Neck Muscles in the rhesus monkey. I. Muscle morphometry and histochemistry. J. Neurophysiol. 86, 1717–1728.

Ruff, C.B., 2003. Long bone articular and diaphyseal structure in Old World Monkeys and Apes. II: Estimation of body mass. Am. J. Phys. Anthropol. 120, 16–37.

Sears, K.E., Bianchi, C., Powers, L., Beck, A.L., 2013. Integration of the mammalian shoulder girdle within populations and over evolutionary time. J. Evol. Biol. 26, 1536– 1548.

Sears, K.E., Capellini, T.S., Diogo, R., 2015. On the serial homology of the pectoral and pelvic girdles of tetrapods. Evolution 69, 2543–2555.

Sigmon, B.A., 1975. Functions and evolution of hominid hip and thigh musculature. In: Tuttle, R. (Ed.), Primate Functional Morphology and Evolution. The Hague, Paris, pp. 235–262.

Stern, J.T., Susman, R.L., 1983. The locomotor anatomy of *Australopithecus afarensis*. Evol. Anthropol. 9, 113–133.

Twede, D.R., Hayden, A.F., 2004. Refinement and generalization of the extension method of covariance matrix inversion by regularizing for spectral filtering optimization. Proc. SPIE 5159, 299–306.

Valasek, P., Theis, S., Krejci, E., Grim, M., Maina, F., Shwartz, Y., Otto, A., Huang, R., Patel, K., 2010. Somatic origin of the medial border of the mammalian scapula and its homology to the avian scapula blade. J. Anat. 216, 482–488.

White, T.D., Black, M.T., Folkens, P.A., 2012. Human Osteology. Third Edition. Academic Press, San Diego.

Young, N.M., 2006. Function, ontogeny and canalization of shape variance in the primate scapula. J. Anat. 209, 623–636.

Zihlman, A., Underwood, C., 2019. Ape Anatomy and Evolution. Independently Published.
